# Genome-scale dissection of phase-variable gene function in *Campylobacter jejuni* using a stabilized phasotype library

**DOI:** 10.64898/2025.12.14.694251

**Authors:** Shouji Yamamoto, Ken-ichi Lee, Akiko Kubomura, Sunao Iyoda, Yukihiro Akeda, Takaaki Shimohata, Chihiro Aikawa, Masashi Okamura, Fuhito Hojo, Takako Osaki, Jiro Mitobe

**Author notes:** Shouji Yamamoto **Email:**.

## Abstract

Phase variation (PV) enables bacterial pathogens to rapidly alter their surface structures through reversible mutations in simple sequence repeats, promoting immune evasion and environmental adaptation. In *Campylobacter jejuni*, the stochastic nature of PV has hindered the systematic functional analysis of phase-variable genes (PVGs). Here, we introduce PV-GenShift, a genome-scale screening platform built on a genetically stabilized library of phase-locked *C. jejuni* variants. By fixing the ON/OFF states of 15 PVGs, PV-GenShift enables reproducible, high-resolution analysis of phasotypes, defined as unique ON/OFF combinations across multiple PVGs, under defined selective pressures. Using models of human serum exposure, murine colonization, and chicken gut passage, we identified distinct phasotypes associated with serum resistance and with enrichment during mouse colonization, particularly involving capsular polysaccharide modifications such as *O*-methyl phosphoramidation and methylation. In contrast, chicken gut passage resulted in heterogeneous ON/OFF shifts without a dominant phasotype. These findings highlight the combinatorial impact of PVG expression states on bacterial adaptation and establish PV-GenShift as a broadly applicable framework for dissecting PV-driven phenotypic diversity. This approach provides a scalable strategy for exploring genotype-phenotype relationships and offers insights relevant to vaccine design and targeted therapeutics.

**Significance Statement:** Phase variation generates phenotypic diversity that enables pathogens to evade immunity and adapt to changing environments; however, its random nature has long obscured functional analysis. This study introduces PV-GenShift, a genome-scale platform that stabilizes phase-variable gene expression in *Campylobacter jejuni*, allowing the systematic identification of gene combinations that influence survival under selective pressures. Using PV-GenShift, we identified phasotypes associated with serum resistance and enrichment during mouse colonization, while chicken passage produced diverse but non-specific shifts. These results demonstrate how combinatorial ON/OFF states of multiple genes shape bacterial adaptation and provide a generalizable strategy for studying phase variation across pathogens, with implications for vaccine design and targeted therapeutics.

## Introduction

Bacterial pathogens have developed sophisticated strategies to survive and adapt in dynamic environments. One such strategy is phase variation (PV), a reversible, high-frequency mechanism of gene regulation that enables clonal populations to generate phenotypic diversity without altering their core genome (1, 2). This stochastic switching of gene expression helps bacteria evade host immune responses (3–6), contributes to pathogenicity (7, 8), facilitates adaptation to environmental pressures (9), and provides resistance against threats such as antibiotics and bacteriophages (10–12). PV typically affects surface-exposed structures, including pili, flagella, outer membrane proteins, and polysaccharides—key components in host-pathogen interactions (1, 13–15).

A major driver of PV is the presence of simple sequence repeats (SSRs), such as polyG/C and polyA/T tracts, which are prone to slipped-strand mispairing during DNA replication. This leads to insertions or deletions that can disrupt coding sequences or regulatory regions, resulting in ON/OFF switching or modulation of gene expression (16, 17) (Fig. 1). SSR-mediated PV has been widely documented in species such as *Neisseria* spp., *Haemophilus influenzae*, *Helicobacter pylori*, and *Campylobacter jejuni* (18–22). Other mechanisms also contribute to PV, including site-specific DNA recombination, such as promoter inversion in *Salmonella enterica* (23, 24), and epigenetic regulation, where phase-variable DNA methylation alters the expression of multiple genes, forming so-called “phasevarions” (25).

**Figure 1.**
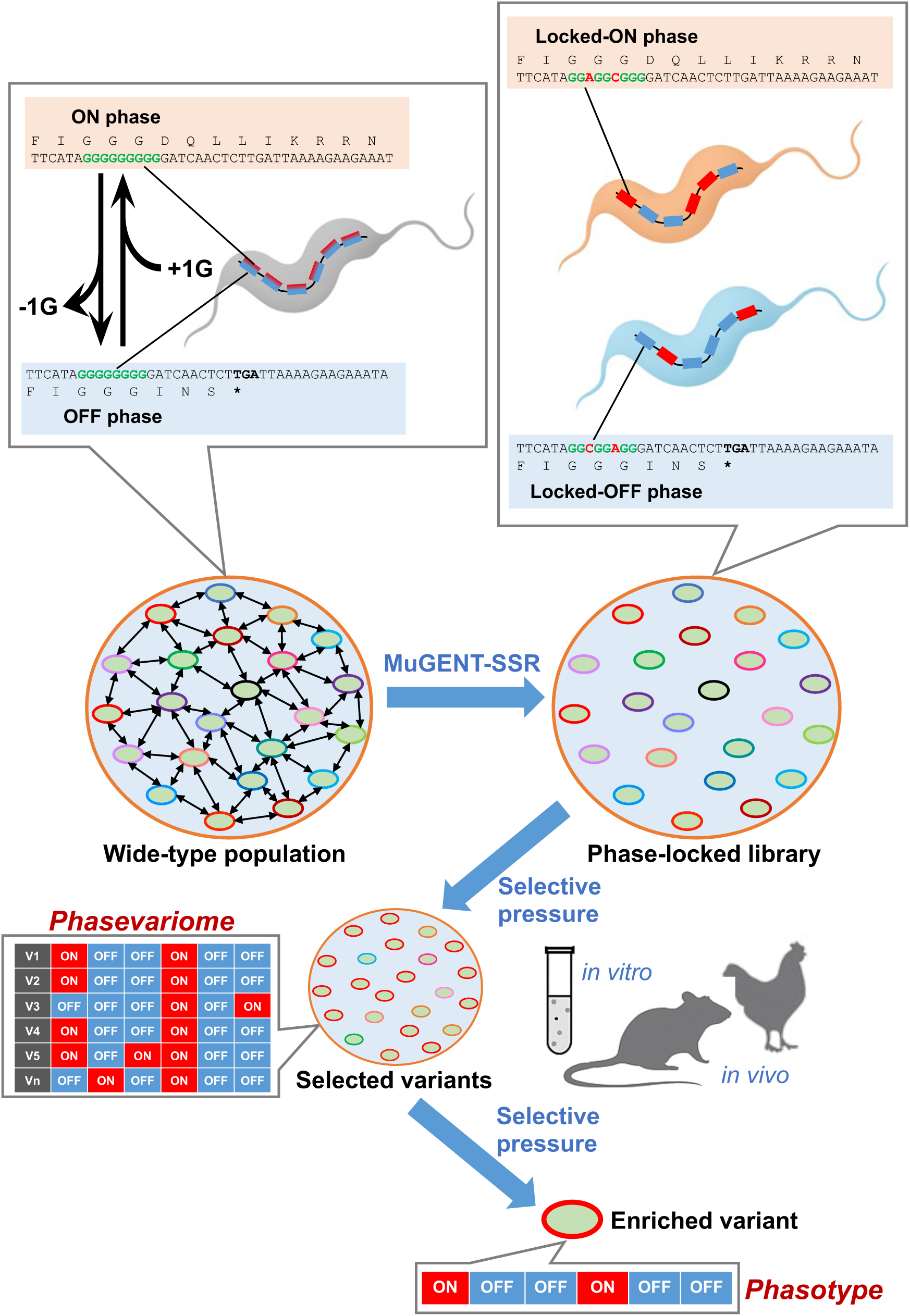
Schematic overview of the PV-GenShift strategy and phasevariome-guided evolutionary screening. PV-GenShift is a stepwise screening framework that utilizes a genetically stabilized phase-locked library of *Campylobacter jejuni* variants. Phase-variable genes (PVGs) containing polyG/C tracts were locked in either the ON or OFF phase using MuGENT-SSR, eliminating stochastic switching and enabling the controlled analysis of phasotypes. In this schematic, six PVGs are shown as an illustrative example, with ON-phase genes indicated in red and OFF-phase genes in blue. This configuration allows for the theoretical generation of 2⁶ (64) distinct phasotypes, but does not represent the full diversity of the phase-locked library. Unlike wild-type populations, which are genetically and phenotypically unstable due to spontaneous phase variation, the phase-locked library provides a highly stable platform for investigating the functional roles of specific phasotypes under defined selective pressures. The phase-locked library was subjected to biologically relevant selective pressures both *in vitro* and *in vivo* (in this study, human serum exposure, colonization in IL-10-knockout mice, and chicken gut passage). Phasevariome analysis of recovered populations revealed distinct ON/OFF expression patterns across PVGs, enabling identification of enriched phasotypes. Once a dominant phasotype is identified, functional validation is performed by switching individual ON-phase genes to the OFF phase to assess their contribution to the observed phenotype. This allows precise identification of causal PVGs involved in host adaptation.

*C. jejuni,* a leading cause of bacterial gastroenteritis worldwide, is also associated with Guillain–Barré syndrome (GBS), a neurological disorder that can occur as a post-infectious complication. Clinical outcomes of *C. jejuni* infection range from mild, watery diarrhea to severe mucoid, bloody diarrhea, reflecting substantial phenotypic diversity (26). The factors determining this diversity are not fully understood; however, it is thought that both extensive genomic variability among strains and heterogeneity driven by PV may have substantial contributions. These two sources of variation complicate efforts to identify stable virulence determinants and predict pathogenic outcomes. Therefore, integrating PV research with genomic studies is essential for understanding host-pathogen interactions and guiding the development of effective interventions. In *C. jejuni*, SSR-mediated PV regulates genes involved in the biosynthesis and modification of lipooligosaccharides (LOS), capsular polysaccharides (CPS), and flagella (27–35)—structures that influence colonization, immune evasion, and virulence (36–44). Notably, PV of LOS biosynthesis genes can lead to molecular mimicry of host gangliosides, contributing to the development of GBS (32). The genetic diversity and phase-variable nature of these surface structures complicate vaccine design (33, 45, 46).

Recent studies show that not only individual phase-variable genes (PVGs), but also their combinatorial ON/OFF states—known as phasotypes (47)—significantly influence bacterial phenotypes, such as serum resistance and phage susceptibility (10, 28, 48). These combinatorial effects suggest that bacterial behavior is shaped by the collective expression states of multiple PVGs, and that interactions among them may further modulate phenotypic outcomes.

In animal models, host passage of *C. jejuni* induces changes in the phasevariome—the complete set of PVGs and their current expression states within a population (47, 49–51). This dynamic PV-driven diversity plays a pivotal role in host adaptation and population heterogeneity. However, the stochastic nature of PV and the complexity of PVG interactions pose major challenges to systematic investigation. Wild-type strains used in previous studies are genetically unstable, undermining reproducibility and obscuring genotype-phenotype correlations. Moreover, *Campylobacter* genomes harbor numerous hypermutable SSRs—typically 18–39 per strain—further complicating efforts to achieve consistent results (52).

To address these limitations, we developed a genome-scale engineering strategy that randomly locks all polyG/C-containing PVGs into defined ON or OFF states, generating a genetically stable and diverse library of phase variants. Building on this, we established the Phase Variation Genomic Shift (PV-GenShift) platform—an experimental evolution framework designed to identify adaptive phasotypes under defined selective pressures (Fig. 1). PV-GenShift enables the systematic dissection of PV dynamics and provides a powerful tool to investigate host-driven selection, vaccine responsiveness, and microbial adaptation.

In this study, we apply PV-GenShift to *C. jejuni* to explore the functional consequences of PV at the genome scale. Our findings offer new insights into the role of PV in bacterial pathogenesis and present a broadly applicable approach for studying phase variation across diverse microbial species.

## Results

### Experimental design of PV-GenShift

PV-GenShift is a stepwise evolutionary screening strategy based on a genetically stabilized library of *C. jejuni* phase-locked variants. By locking the ON/OFF states of PVGs, this approach enables systematic dissection of their roles in host adaptation under defined selective pressures (Fig. 1).

#### 1) Library Construction and Validation

Using multiplex genome editing, we selectively targeted polyG/C-containing SSRs within coding regions, excluding promoter and Shine–Dalgarno (SD) sequences, to randomly lock each PVG in either an ON or OFF state. This generated a diverse library of variants with defined and stable phasotypes. Whole-genome sequencing (WGS) confirmed precise editing with no off-target mutations, and the engineered strains maintained consistent gene expression profiles across passages.

#### 2) Stepwise Screening Under Selective Pressures

To demonstrate the versatility of the PV-GenShift platform, we applied the phase-locked variant library to a series of biologically relevant conditions that simulate host environments. In this study, we selected three representative models as illustrative examples: exposure to human serum to assess complement resistance, colonization in mice to evaluate intestinal fitness, and chicken gut passage to investigate host-specific colonization potential. These selective pressures were applied in a stepwise manner, allowing for the simultaneous comparison of multiple phasotypes and the enrichment of variants with enhanced survival or colonization traits. While these models were chosen as examples for this study, the PV-GenShift framework may be broadly applicable to a wide range of selective environments, such as antimicrobial exposure, phage challenge, or nutrient limitation.

#### 3) Phasevariome Analysis

WGS of recovered populations enabled characterization of the phasevariome—the complete set of PVG expression states within each population. While wild-type *C. jejuni* exhibits a highly dynamic phasevariome due to stochastic switching, our stabilized variants allowed reproducible analysis of its role in adaptation.

#### 4) Phasotype and Gene-Level Dissection

Dominant phasotypes enriched under selection were identified, and ON-phase genes within these phasotypes were analyzed to determine specific PVGs contributing to the emergence of adaptive traits. This enabled assessment of both individual and combinatorial effects of PVG expression on host interaction.

By stabilizing PVG states and applying stepwise evolutionary screening, PV-GenShift allows isolation of phenotypes that are typically transient or masked by population heterogeneity. This platform provides new insights into *C. jejuni* pathogenesis and facilitates a detailed exploration of genotype-phenotype relationships through the lens of the phasevariome.

### Construction and evaluation of a phase-locked variant library in *C. jejuni* strain 81-176

To enable a systematic analysis of PVG function, we constructed and evaluated a genetically stabilized library of phase-locked variants in *C. jejuni* strain 81-176. This strain contains 18 polyG/C SSR tracts, 15 of which are located within open reading frames (ORFs) and are associated with PVG expression (52) (Table 1). In this study, the remaining three tracts located in intergenic regions were excluded, as the potential impact of SSR repeat variation on gene expression remains unclear. Among the 15 PVGs analyzed, we considered *CJJ81176_1421* as a single PVG, despite its overlapping annotation with *CJJ81176_1420*. This decision was based on the presence of a clear SD sequence upstream of *CJJ81176_1421* and the absence of an independent SD sequence for *CJJ81176_1420*, as well as the shared reading frame between the two (Fig. S1). These observations suggest that *CJJ81176_1420* is likely a misannotation and that the region represents a single functional PVG, *CJJ81176_1421*, regulated by SSR-mediated PV.

**Table 1.**
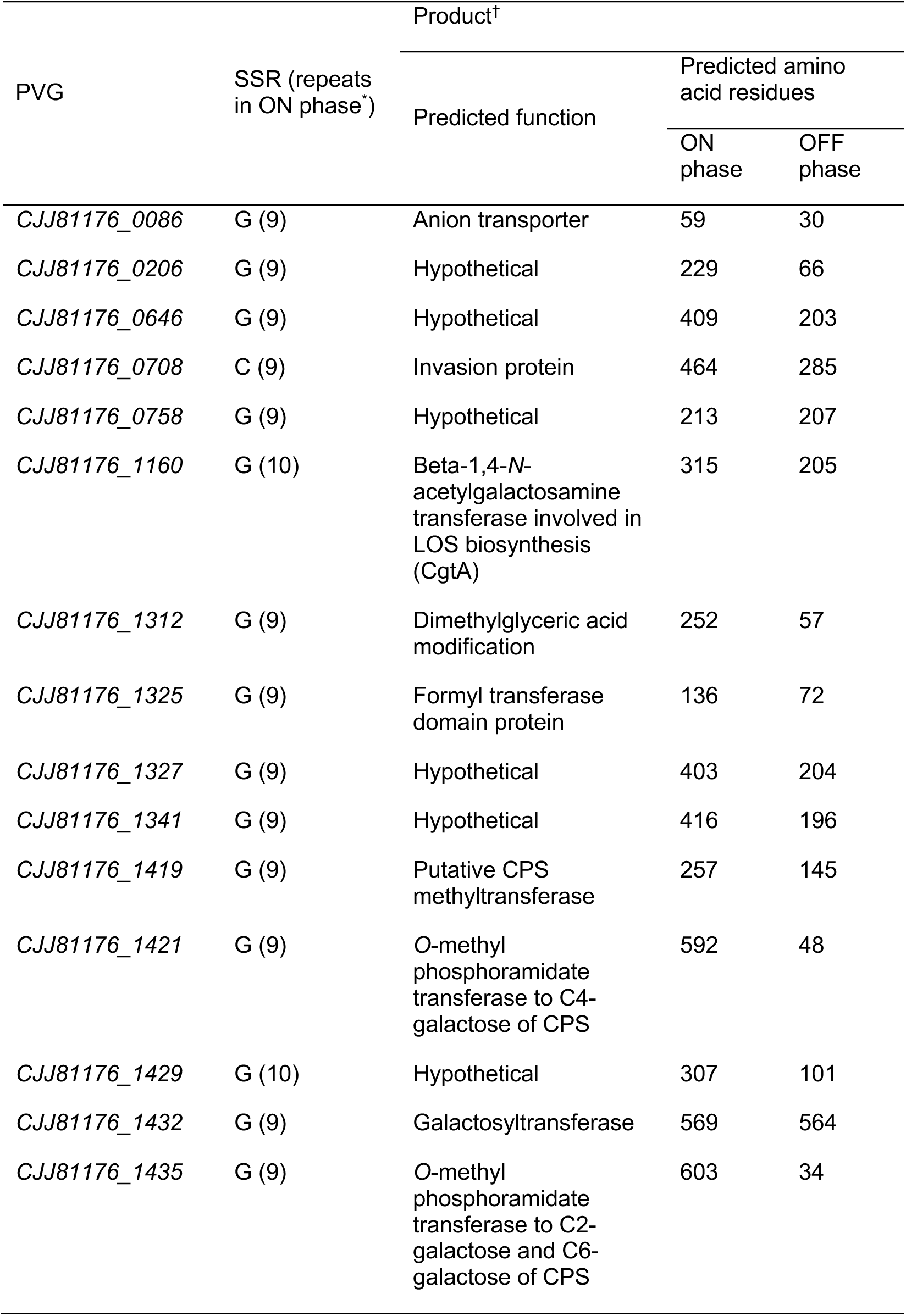
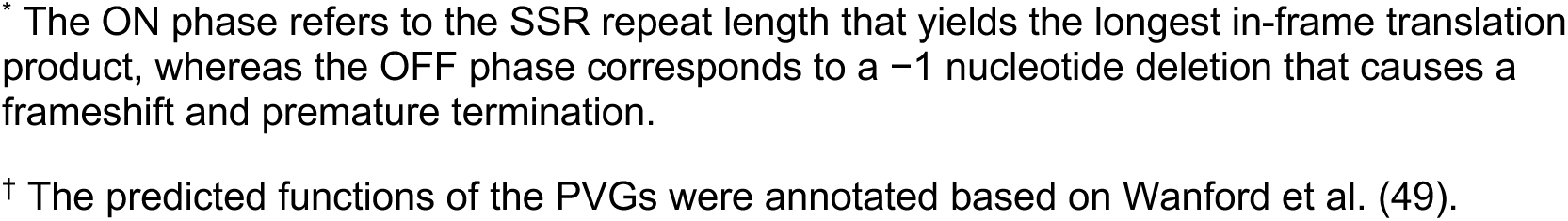
Predicted products of polyG/C-containing PVGs stabilized via MuGENT-SSR in *Campylobacter jejuni* 81-176.

| PVG | SSR (repeats<br>in ON phase*) | Product<sup>†</sup> |  |  |
| --- | --- | --- | --- | --- |
|  |  | Predicted function | Predicted amino<br>acid residues |  |
|  |  |  | ON<br>phase | OFF<br>phase |
| *CJJ81176_0086* | G (9) | Anion transporter | 59 | 30 |
| *CJJ81176_0206* | G (9) | Hypothetical | 229 | 66 |
| *CJJ81176_0646* | G (9) | Hypothetical | 409 | 203 |
| *CJJ81176_0708* | C (9) | Invasion protein | 464 | 285 |
| *CJJ81176_0758* | G (9) | Hypothetical | 213 | 207 |
| *CJJ81176_1160* | G (10) | Beta-1,4-*N*-<br>acetylgalactosamine<br>transferase involved in<br>LOS biosynthesis<br>(CgtA) | 315 | 205 |
| *CJJ81176_1312* | G (9) | Dimethylglyceric acid<br>modification | 252 | 57 |
| *CJJ81176_1325* | G (9) | Formyl transferase<br>domain protein | 136 | 72 |
| *CJJ81176_1327* | G (9) | Hypothetical | 403 | 204 |
| *CJJ81176_1341* | G (9) | Hypothetical | 416 | 196 |
| *CJJ81176_1419* | G (9) | Putative CPS<br>methyltransferase | 257 | 145 |
| *CJJ81176_1421* | G (9) | O-methyl<br>phosphoramidate<br>transferase to C4-galactose of CPS | 592 | 48 |
| *CJJ81176_1429* | G (10) | Hypothetical | 307 | 101 |
| *CJJ81176_1432* | G (9) | Galactosyltransferase | 569 | 564 |
| *CJJ81176_1435* | G (9) | O-methyl<br>phosphoramidate<br>transferase to C2-galactose and C6-galactose of CPS | 603 | 34 |

\* The ON phase refers to the SSR repeat length that yields the longest in-frame translation product, whereas the OFF phase corresponds to a -1 nucleotide deletion that causes a frameshift and premature termination.
† The predicted functions of the PVGs were annotated based on Wanford et al. (49).

SSRs within ORFs are prone to slipped-strand mispairing during DNA replication, leading to reversible ON/OFF switching via frameshift mutations. For this study, the ON phase was defined as the SSR length that produces the longest in-frame translation product, while the OFF phase was defined as a −1 nucleotide deletion that induces a frameshift and premature termination (Table 1). This mechanism theoretically allows for up to 2¹⁵ distinct phasotypes per genome.

To generate a genetically stable and fully edited population of phasotypes, we employed MuGENT-SSR (Multiplex Genome Editing by Natural Transformation against Simple Sequence Repeats), a high-efficiency genome editing method based on natural transformation and co-transformation (53) (Fig. 2). This approach utilizes two types of donor DNA: (i) selected DNA containing an antibiotic resistance marker (kanamycin or chloramphenicol) inserted into a neutral locus (e.g., *flaA*), and (ii) unselected DNA comprising 15 fragments designed to introduce mutations into the polyG/C tracts of PVGs, thereby locking each gene in either the ON or OFF phase.

**Figure 2.**
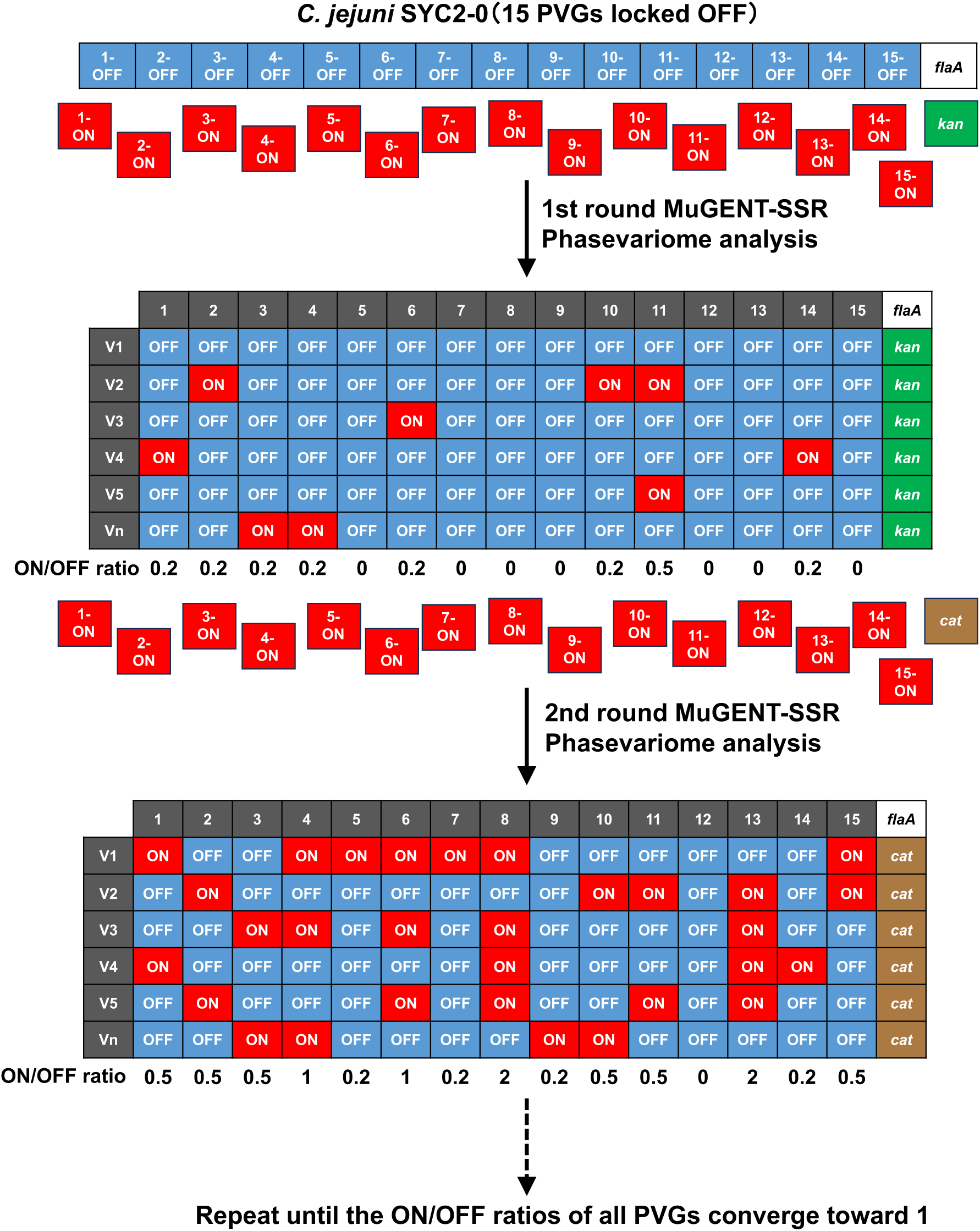
Construction and evaluation of a phase-locked variant library in *Campylobacter jejuni* strain 81-176. In the first round of MuGENT-SSR, the *C. jejuni* 81-176 variant SYC2-0, in which all 15 phase-variable genes (PVGs) were locked in the OFF phase, was used as the starting strain. A total of 15 PVGs containing polyG/C tracts within coding regions were genetically stabilized using MuGENT-SSR. Each PVG was locked in either the ON or OFF phase, resulting in a diverse library of phase-locked variants. The ON phase corresponds to a simple sequence repeat (SSR) length that maintains the correct reading frame for full-length protein translation, whereas the OFF phase is induced by a −1 nucleotide frameshift, leading to premature termination. Phasotypes of variants (V1–Vn) are shown as illustrative examples, with ON-phase genes indicated in red and OFF-phase genes in blue. These examples highlight the combinatorial diversity of PVG expression states across the library. MuGENT-SSR was performed in sequential rounds using selected DNA fragments carrying antibiotic resistance markers (kanamycin or chloramphenicol) targeted to *flaA*, along with unselected DNA fragments designed to lock PVGs in the ON state. Whole-genome sequencing of pooled colonies (20,000–30,000 per round) was used to assess editing efficiency and phase distribution. ON/OFF ratios for each PVG were calculated using PVfinder_81176, a BLAST-based computational tool developed for this study. The ON/OFF ratio shown below the table was calculated from six data points (V1 through Vn) as an illustrative example. The editing process was repeated until the ON/OFF ratios of all PVGs converged toward 1, indicating a balanced and complete phasevariome suitable for downstream screening.

To maintain transformation efficiency and avoid repeated selection at the same locus, a different DNA construct carrying a distinct antibiotic resistance marker was used in each MuGENT cycle. Through sequential rounds, we progressively increased the incorporation of locked ON or OFF mutations across the population, enhancing mutation coverage and ensuring that all targeted PVGs were successfully edited (Fig. 2).

In the initial step, all 15 PVGs were locked in the OFF phase by co-transforming cells with selected DNA and unselected fragments carrying locked-OFF mutations (e.g., GGG GGG GGG → GGA GGC GG), preventing slipped-strand mispairing and maintaining the gene in the OFF state. Antibiotic selection yielded a population in which all PVGs were stably locked OFF. This population was then subjected to a second round of MuGENT-SSR using unselected DNA fragments carrying locked-ON mutations (e.g., GGG GGG GGG → GGC GGA GGG), randomly converting a subset of PVGs back to the ON phase. Repeating this ON-switching step with different combinations of unselected DNA further increased the number of edited loci, generating a collection of phase-locked variants with stable and combinatorially distinct phasotypes (Fig. 2).

To quantitatively assess the completeness and balance of PVG editing, we performed short-read WGS on pooled genomic DNA (gDNA) extracted from approximately 20,000 to 30,000 colonies obtained after MuGENT-SSR. Sequencing data were then analyzed using PVfinder_81176, a BLAST-based computational tool developed for this study. PVfinder_81176 calculates the ON/OFF ratio for each PVG by comparing the number of reads corresponding to locked-ON and locked-OFF alleles (Table S5). An ON/OFF ratio close to 1 indicates balanced representation of both phases, while ratios approaching ∞ or 0 indicate dominance of ON or OFF variants, respectively. The ON/OFF ratio profile across all PVGs—termed the phasevariome—was used to evaluate and select libraries with complete and well-distributed phasotype compositions (Fig. 2).

To further improve mutation coverage, we performed five iterative rounds of MuGENT-SSR, each aimed at randomly switching previously locked-OFF PVGs to the ON state (Table 2 and Table S6). At each round, the phasevariome was analyzed using PVfinder_81176. While all PVGs exhibited changes in their ON/OFF ratios over time, the rate of change varied considerably. Some genes reached a near-equilibrium ratio of approximately 1 by the third round, indicating balanced representation of ON and OFF alleles. In contrast, others remained strongly biased toward the OFF phase, with ratios below 0.3 even after the fifth round.

**Table 2.** Phasevariome profiles during the construction of the phase-locked library in

| PVG | ON/OFF Ratios* of PVGs Across MuGENT-SSR Round |  |  |  |  |  |
| --- | --- | --- | --- | --- | --- | --- |
|  | 0 | 1 | 2 | 3 | 4 | 5 |
| *CJJ81176_0086* | 0 | 0.10 | 0.12 | 0.17 | 0.23 | 0.27 |
| *CJJ81176_0206* | 0 | 0.20 | 0.30 | 0.34 | 0.40 | 0.44 |
| *CJJ81176_0646* | 0 | 0.08 | 0.12 | 0.16 | 0.13 | 0.27 |
| *CJJ81176_0708* | 0 | 0.17 | 0.18 | 0.25 | 0.33 | 0.38 |
| *CJJ81176_0758* | 0 | 0.09 | 0.15 | 0.17 | 0.21 | 0.32 |
| *CJJ81176_1160* | 0 | 0.24 | 0.31 | 0.91 | 1.01 | 1.50 |
| *CJJ81176_1312* | 0 | 0.11 | 0.10 | 0.12 | 0.19 | 0.35 |
| *CJJ81176_1325* | 0 | 0.20 | 0.21 | 0.29 | 0.55 | 0.41 |
| *CJJ81176_1327* | 0 | 0.14 | 0.19 | 0.34 | 0.40 | 0.52 |
| *CJJ81176_1341* | 0 | 0.06 | 0.12 | 0.15 | 0.26 | 0.23 |
| *CJJ81176_1419* | 0 | 0.15 | 0.14 | 0.69 | 0.62 | 1.08 |
| *CJJ81176_1421* | 0 | 0.05 | 0.02 | 0.07 | 0.15 | 0.23 |
| *CJJ81176_1429* | 0 | 0.33 | 0.48 | 0.74 | 1.97 | 0.93 |
| *CJJ81176_1432* | 0 | 0.27 | 0.32 | 0.51 | 0.53 | 0.31 |
| *CJJ81176_1435* | 0 | 0.08 | 0.10 | 0.56 | 0.36 | 0.91 |

\* PVfinder\_81176 calculated the ON/OFF ratio for each PVG by comparing the number of reads corresponding to the locked-ON and locked-OFF alleles.

Given the constraints of the experimental timeline, we selected the bacterial population from the fifth round as the final phase-locked variant library, designated PLL_81176. To restore motility, which had been disrupted by the insertion of an antibiotic resistance cassette into the *flaA* locus, we repaired the *flaA* gene to its wild-type sequence via natural transformation using unmarked donor DNA. This step successfully reconstituted functional flagella, enabling motile behavior. As a result, we established a genetically stable and motile phase-locked variant library, designated PLL_81176M, consisting of approximately 20,000 independent colonies. This population is expected to encompass a wide range of combinatorially distinct phasotypes, providing a robust resource for downstream phenotypic screening and evolutionary analyses.

### Evolutionary screening of human serum-resistant variants from PLL_81176M and identification of causative genes

*C. jejuni* is known for its resistance to human serum, antimicrobial peptides, and bacteriophages. Surface antigens such as CPS and LOS play critical roles in these defense mechanisms (10, 28, 38, 39, 54). These structures also facilitate host cell invasion (29, 38, 55) and are associated with various diseases, including diarrhea and GBS (39, 41, 42, 56). Since PV drives the structural diversity of these antigens, understanding how PV influences host interactions and immune evasion is essential.

To evaluate the utility of the PV-GenShift platform, we conducted evolutionary screening and genome-wide phasevariome analysis using serum resistance as a model phenotype. Briefly, cultures of PLL_81176M (input) were exposed to human serum, and surviving colonies (output) were collected and re-exposed in subsequent rounds. Consistent with previous studies, no colonies survived serum exposure in the *kpsE* deletion mutant, which lacks CPS production (55) (Table S1, SYC2005). The serum resistance rate (calculated as 100 × output CFU/input CFU) increased progressively with each round of exposure, reaching a peak after the third round. By this point, the resistance rate was 33-fold higher than the initial exposure, indicating that repeated serum exposure enriches for variants with enhanced resistance. In contrast, the parental strain 81-176 showed a resistance rate of only 13.5 ± 4.6% after a single exposure.

Phasevariome analysis of input and output samples revealed consistent ON/OFF shifts in several PVGs. Specifically, *CJJ81176_0758*, *CJJ81176_1419*, *CJJ81176_1421*, and *CJJ81176_1435* shifted more than 10-fold toward the ON phase, while *CJJ81176_1429* shifted more than 10-fold toward the OFF phase (Table 3 and Table S7). To further investigate the genetic basis of resistance, 5 single colonies from the third output were isolated and assigned phasotypes (PTs), which are numerical identifiers based on the binary ON/OFF status of PVGs. Three distinct PTs were identified: PT1211, PT4263, and PT9397, with PT9397 showing the highest serum resistance (Table 4).

**Table 3.** Phasevariome of human serum-resistant variants from PLL_81176.

| PVG | ON/OFF Ratios of PVGs |  |  |  |
| --- | --- | --- | --- | --- |
|  | Input | Output |  |  |
|  |  | 1st | 2nd | 3rd |
| *CJJ81176_0086* | 0.25 | 0.17 | 0.12 | 0.10 |
| *CJJ81176_0206* | 0.32 | 0.32 | 0.35 | 0.42 |
| *CJJ81176_0646* | 0.31 | 0.39 | 0.15 | 0.09 |
| *CJJ81176_0708* | 0.29 | 0.38 | 0.12 | 0.06 |
| *CJJ81176_0758* | 0.30 | 1.44 | 6.53 | 7.98 |
| *CJJ81176_1160* | 0.06 | 0.17 | 0.12 | 0.06 |
| *CJJ81176_1312* | 0.23 | 0.22 | 0.10 | 0.09 |
| *CJJ81176_1325* | 24.13 | 8.47 | 15.79 | 23.63 |
| *CJJ81176_1327* | 0.19 | 0.38 | 0.09 | 0.08 |
| *CJJ81176_1341* | 27.15 | 7.72 | 10.94 | 22.63 |
| *CJJ81176_1419* | 0.06 | 4.47 | 24.62 | 28.04 |
| *CJJ81176_1421* | 0.08 | 3.21 | 4.33 | 2.73 |
| *CJJ81176_1429* | 7.01 | 0.46 | 0.34 | 0.41 |
| *CJJ81176_1432* | 0.36 | 0.82 | 1.43 | 1.49 |
| *CJJ81176_1435* | 0.13 | 1.92 | 10.56 | 56.50 |
| Serum Resistance Rate ± SD (%)* | NA | 11.0 ± 2.6 | 122.3 ± 19.1 | 120.3 ± 7.8 |

\* Values indicate the serum resistance rate ± standard deviation (SD). The serum resistance rate (%) was calculated as: (number of surviving output bacteria after serum exposure ÷ number of input bacteria before exposure) × 100. The assay was performed in triplicate. NA indicates not applicable.

**Table 4.** Identification of phasotypes and causative genes involved in serum resistance in

| PVG | Phasotype* |  |  |  |  |  |  |  |  |  |
| --- | --- | --- | --- | --- | --- | --- | --- | --- | --- | --- |
|  | #1 | #2 | #3 | #4 | #5 | #3v1 | #3v2 | #3v3 | #3v4 | #3v5 |
| CJJ81176_0086 | 0 | 0 | 0 | 0 | 0 | 0 | 0 | 0 | 0 | 0 |
| CJJ81176_0206 | 0 | 0 | 1 | 1 | 0 | 0 | 0 | 0 | 0 | 0 |
| CJJ81176_0646 | 1 | 1 | 0 | 0 | 0 | 0 | 0 | 0 | 0 | 0 |
| CJJ81176_0708 | 0 | 0 | 0 | 0 | 0 | 0 | 0 | 0 | 0 | 0 |
| CJJ81176_0758 | 0 | 1 | 1 | 1 | 1 | 0 | 0 | 0 | 0 | 0 |
| CJJ81176_1160 | 0 | 0 | 0 | 0 | 0 | 0 | 0 | 0 | 0 | 0 |
| CJJ81176_1312 | 0 | 0 | 0 | 0 | 0 | 0 | 0 | 0 | 0 | 0 |
| CJJ81176_1325 | 1 | 1 | 1 | 1 | 1 | 0 | 0 | 0 | 0 | 0 |
| CJJ81176_1327 | 0 | 0 | 0 | 0 | 0 | 0 | 0 | 0 | 0 | 0 |
| CJJ81176_1341 | 1 | 1 | 1 | 1 | 1 | 0 | 0 | 0 | 0 | 0 |
| CJJ81176_1419 | 0 | 1 | 1 | 1 | 1 | 1 | 0 | 1 | 1 | 0 |
| CJJ81176_1421 | 0 | 1 | 0 | 0 | 1 | 0 | 0 | 0 | 0 | 0 |
| CJJ81176_1429 | 1 | 0 | 1 | 1 | 0 | 1 | 1 | 0 | 1 | 0 |
| CJJ81176_1432 | 1 | 1 | 0 | 0 | 1 | 0 | 0 | 0 | 0 | 0 |
| CJJ81176_1435 | 1 | 1 | 1 | 1 | 1 | 1 | 1 | 1 | 0 | 0 |
|  | 4263 | 1211 | 9397 | 9397 | 1211 | 21 | 5 | 17 | 20 | 0 |
| Serum Resistance<br>Rate ± SD (%)<sup>†</sup> | 31 ± 4 | 78 ± 5 | 103 ± 16 | 114 ± 8 | 85 ± 16 | 96 ± 11 | ND | ND | ND | ND |

\* Five single colonies (#1 to #5) were isolated from the output of the third exposure, and their genome sequences were analyzed to determine their phasotypes and to conduct serum resistance assays. The phase status of each gene is represented in binary format, with ON as 1 and OFF as 0. Additionally, the phasotypes were converted from binary to decimal format as shown in the bottom row. For example, converting the binary 001000010100111 to decimal gives 4263. Based on the phasotype 9397 (#3), various mutant strains (#3v1 to #3v5) were constructed and subjected to serum resistance testing. The following strains were used: #3, SYC2-SV1; #3v1, SYC2-SV2; #3v2, SYC2-SV3; #3v3, SYC2-SV4; #3v4, SYC2-SV5; and #3v5, SYC2-SV6.
† ND indicates that surviving output colonies were not detected.

To identify causative genes, ON-phase genes in PT9397 were individually switched to the OFF state. Serum resistance was retained only when *CJJ81176_1419*, *CJJ81176_1429*, and *CJJ81176_1435* remained ON (PT21) (Table 4). Among these, *CJJ81176_1435* encodes an *O*-methyl phosphoramidate (MeOPN) transferase responsible for adding MeOPN to both the C2 and C6 positions of galactose of the 81-176 CPS and is linked to serum resistance (28). *CJJ81176_1419* encodes a homolog of *cj1420*, a putative methyltransferase involved in CPS biosynthesis in *C. jejuni* NCTC11168, which may influence both serum and phage sensitivity (49). *CJJ81176_1429* is a conserved gene of unknown function.

### Evolutionary screening of PLL_81176M colonization in IL-10-knockout mice

The previous *in vitro* results demonstrated that the PV-GenShift platform can effectively enrich serum-resistant variants and identify both known and novel PVGs contributing to immune resistance in *C. jejuni*. To explore whether specific phasotypes are selectively enriched during host colonization, we applied PV-GenShift in an *in vivo* setting using the PLL_81176M library.

Previous studies have shown that C57BL/6 IL-10⁺/⁺ mice are suitable for colonization studies, while IL-10⁻/⁻ mice can serve as a model for both colonization and enteritis (57). In a preliminary experiment, we administered PLL_81176M (2.8 × 10⁹ CFU) or its parent strain 81-176 (1.0 × 10¹⁰ CFU) to three IL-10⁺/⁺ mice. After 7 days, no bacterial colonies were detected in fecal samples, indicating a failure to colonize under non-inflammatory conditions. In contrast, IL-10⁻/⁻ mice infected with 81-176 showed successful colonization, with fecal bacterial loads ranging from 2.7 × 10⁴ to 6.8 × 10⁶ CFU/g.

Based on these observations, we conducted PV-GenShift screening using PLL_81176M in IL-10⁻/⁻ mice to investigate the *in vivo* relevance of specific phasotypes under inflammatory conditions. Briefly, a suspension of 3.0 × 10⁸ CFU was administered to three IL-10⁻/⁻ mice. Seven days post-infection, clinical symptoms were assessed, and fecal and cecal samples were collected. Bacterial loads ranged from 1.1 × 10⁸ to 6.7 × 10⁸ CFU/g in the feces and from 9.0 × 10⁸ to 1.1 × 10⁹ CFU/g in the cecal contents, confirming successful colonization (Table 5). Macroscopic examination of the colons from IL-10⁻/⁻ mice revealed mucosal erythema, edema, and marked wall thickening. However, it remains unclear whether these changes were directly caused by infection or were a consequence of IL-10 deficiency.

**Table 5.** Phasevariome of PLL_81176-derived colonizing variants in the cecum and feces of IL-10-knockout mice.

| PVG | ON/OFF Ratios of PVGs |  |  |  |  |  |  |
| --- | --- | --- | --- | --- | --- | --- | --- |
|  | Input | Output* |  |  |  |  |  |
|  |  | m#1 |  | m#2 |  | m#3 |  |
|  |  | Cecum | Feces | Cecum | Feces | Cecum | Feces |
| *CJJ81176_0086* | 0.29 | 0.00 | 0.00 | 0.00 | 0.00 | 0.00 | 0.00 |
| *CJJ81176_0206* | 0.46 | 0.01 | 0.01 | 0.01 | 0.01 | 0.09 | 0.04 |
| *CJJ81176_0646* | 0.53 | 0.00 | 0.01 | 0.00 | 0.01 | 0.01 | 0.01 |
| *CJJ81176_0708* | 0.71 | 0.01 | 0.00 | 0.00 | 0.00 | 0.06 | 0.03 |
| *CJJ81176_0758* | 0.43 | 0.00 | 0.00 | 0.00 | 0.00 | 0.00 | 0.00 |
| *CJJ81176_1160* | 0.42 | 0.00 | 0.01 | 0.00 | 0.01 | 0.01 | 0.02 |
| *CJJ81176_1312* | 0.45 | 268.50 | 690 | 620.00 | 1097 | 12.61 | 42.5 |
| *CJJ81176_1325* | 0.69 | 0.00 | 0.00 | 0.00 | 0.00 | 0.00 | 0.00 |
| *CJJ81176_1327* | 0.67 | 0.00 | 0.00 | 0.00 | 0.01 | 0.01 | 0.01 |
| *CJJ81176_1341* | 0.35 | 0.00 | 0.00 | 0.01 | 0.00 | 0.00 | 0.01 |
| *CJJ81176_1419* | 0.77 | 321.75 | 195 | 639.50 | 1166 | 13.19 | 43.69 |
| *CJJ81176_1421* | 0.49 | 67.67 | 116 | 104.00 | 224 | 21.75 | 51.5 |
| *CJJ81176_1429* | 2.26 | 205.40 | 304.5 | 189.00 | 283.75 | 130.00 | 55.7 |
| *CJJ81176_1432* | 0.94 | 0.00 | 0.00 | 0.00 | 0.00 | 0.00 | 0.01 |
| *CJJ81176_1435* | 0.19 | 0.00 | 0.00 | 0.00 | 0.02 | 0.00 | 0.01 |
| Number of bacteria (CFUs/g) | NA | $1 \times 10^9$ | $6.7 \times 10^8$ | $9 \times 10^8$ | $1.1 \times 10^8$ | $1.1 \times 10^9$ | $2.2 \times 10^8$ |

\* Results of the phasevariome analysis and colony recovery from the cecum contents and feces of three individual mice are shown.

Phasevariome analysis of pooled output colonies revealed consistent ON/OFF shifts across all PVGs. Four genes—*CJJ81176_1312*, *CJJ81176_1419*, *CJJ81176_1421*, and *CJJ81176_1429*—shifted to the ON phase, while the remaining PVGs shifted to the OFF phase (Table 5 and Table S8). The dominant phasotype was PT284, as confirmed in all five single colonies isolated from the output. Three of the ON-phase genes—*CJJ81176_1419*, *CJJ81176_1421*, and *CJJ81176_1429*—are located within the CPS biosynthesis cluster. *CJJ81176_1419* and *CJJ81176_1429* have previously been linked to serum resistance (Table 3). *CJJ81176_1421* encodes an additional MeOPN transferase within the CPS biosynthesis cluster, complementing the previously described *CJJ81176_1435* (Table 3) and is involved in the transfer of MeOPN to the C4 position of galactose of the 81-176 CPS (28). The fourth gene in the ON-phase, *CJJ81176_1312*, is a homolog of *cj1295* in *C. jejuni* NCTC11168. It is predicted to be involved in the addition of dimethylglyceric acid to *O*-linked pseudaminic acid residues found in flagella (31), which may affect bacterial motility and immune evasion.

### Evolutionary screening of PLL_81176M via chicken gut passage

Chickens are a natural reservoir for *C. jejuni* and a major source of human campylobacteriosis through contaminated poultry products. To investigate PV and host adaptation in a biologically relevant host, we applied the PV-GenShift platform to the PLL_81176M library using a chicken infection model.

Herein, a bacterial suspension containing approximately 10⁴ CFU of PLL_81176M (input) was orally administered to individually housed chickens at 27 days of age. Nineteen days post-inoculation, cecal droppings were collected. Bacterial counts recovered from the cecal droppings ranged from 2.3 × 10⁷ to 6.8 × 10^7^ CFU/g (Table 6). Approximately 1,000 to 6,000 colonies from each of three chickens were pooled to generate the output samples.

**Table 6.** Phasevariome of PLL_81176-derived variants recovered from chicken cecal droppings.

| PVG | ON/OFF Ratios of PVGs |  |  |  |
| --- | --- | --- | --- | --- |
|  | Input | Output* |  |  |
|  |  | c#1 | c#2 | c#3 |
| *CJJ81176_0086* | 0.32 | 0.48 | 222.67 | 0.00 |
| *CJJ81176_0206* | 0.49 | 0.00 | 140.88 | 0.00 |
| *CJJ81176_0646* | 0.57 | 1.94 | 695.00 | 0.00 |
| *CJJ81176_0708* | 0.69 | 13.67 | 189.25 | 0.00 |
| *CJJ81176_0758* | 0.54 | 0.00 | 0.00 | ∞ |
| *CJJ81176_1160* | 0.37 | 0.40 | 190.00 | 0 |
| *CJJ81176_1312* | 0.51 | 70.00 | 0.00 | 0 |
| *CJJ81176_1325* | 0.52 | 1.76 | ∞ | ∞ |
| *CJJ81176_1327* | 0.69 | 0.41 | 0.01 | ∞ |
| *CJJ81176_1341* | 0.33 | 0.00 | 0.00 | 0.00 |
| *CJJ81176_1419* | 0.56 | 0.51 | 372.67 | ∞ |
| *CJJ81176_1421* | 0.20 | 0.04 | ∞ | ∞ |
| *CJJ81176_1429* | 2.18 | ∞ | ∞ | ∞ |
| *CJJ81176_1432* | 0.92 | 1.89 | ∞ | 0.00 |
| *CJJ81176_1435* | 0.53 | ∞ | 151.00 | 0.00 |
| Number of bacteria in the cecal droppings (CFUs/g) | NA | $2.3 \times 10^7$ | $4.2 \times 10^7$ | $6.8 \times 10^7$ |

\* The results of the phasevariome analysis and colony recovery from the cecal droppings of three individual chickens are presented.

To assess the impact of host passage on PVG expression, we analyzed the phasevariome of PLL_81176M in chickens before and after infection in chickens. Phasevariome profiles were compared between the input library and the outputs recovered post-infection (Table 6 and Table S9). Even after a single round of infection, all chickens exhibited dramatic shifts in the ON/OFF ratios across PVGs, although the patterns varied among individuals. Notably, *CJJ81176_1429* consistently shifted to the ON phase, while *CJJ81176_1341* shifted to the OFF phase in all chickens. *CJJ81176_1429* has previously shown an involvement in serum resistance (Table 4) and ON-phase shifts following passage in a mouse model (Table 5), suggesting a conserved role in host adaptation. *CJJ81176_1341* encodes a hypothetical protein (Table 1), and its consistent OFF-phase shift may indicate a selective disadvantage during chicken colonization.

## Discussion

PV is a key adaptive strategy in *C. jejuni*, enabling rapid and reversible changes in surface structures through mutations in SSRs. This stochastic switching facilitates immune evasion and environmental adaptation but has historically complicated the functional analysis of individual PVGs. Our findings reinforce the need to consider both genomic diversity and PV-driven heterogeneity when investigating *C. jejuni* pathogenesis. While genome sequencing provides critical insights into strain-specific gene content, PV introduces an additional layer of complexity by enabling reversible ON/OFF switching of multiple loci. Combining genomic approaches with systematic PV analysis is crucial for elucidating mechanisms underlying phenotypic variability and disease severity (58).

In this study, we developed PV-GenShift, a genome-scale evolutionary screening platform based on a genetically stabilized library of phase-locked *C. jejuni* variants. By locking the ON/OFF states of 15 PVGs, PV-GenShift enables a reproducible, high-resolution analysis of phasotype dynamics under defined selective pressures. Using this approach, we identified specific phasotypes that enhance serum resistance, providing direct evidence that distinct PVG configurations contribute to host-associated adaptation (Table 4). In contrast, while certain phasotypes were enriched during host colonization, definitive functional configurations were not identified. Notably, in mouse models, specific phasotypes were consistently enriched post-colonization, suggesting selective pressure for particular PVG states (Table 5). However, in chicken models, phasevariome analysis revealed divergent phasotypes across individuals, with no consistent enrichment (Table 6). This variability may reflect non-selective bottlenecks during colonization, which can obscure phasotype-specific selection and complicate the interpretation of PVG function in avian hosts (49, 59).

In this study, PVGs involved in CPS biosynthesis were consistently enriched in the ON phase across three host-relevant models—human serum exposure, murine colonization, and chicken gut passage. In the serum resistance model, the ON-phase expression of *CJJ81176_1419*, *CJJ81176_1429*, and *CJJ81176_1435* was strongly associated with increased survival. In mice, *CJJ81176_1419*, *CJJ81176_1421*, and *CJJ81176_1429* formed a dominant phasotype. In chickens, an ON-phase shift of *CJJ81176_1429* was observed, although no single phasotype predominated. These genes encode enzymes that modify CPS: CJJ81176_1421 adds MeOPN to the C4 position of galactose, CJJ81176_1435 modifies the C2 and C6 positions (28), and CJJ81176_1419 is predicted to encode a methyltransferase; however, the function of CJJ81176_1429 remains unknown. In strain NCTC11168, combinations of MeOPN expression and methylation have been shown to affect serum resistance, phage susceptibility, and antigenic variation (10). Notably, our previous work demonstrated that Penner serotype antigenicity is determined by the PVG *cj1426*, which is predicted to be involved in CPS methylation, reinforcing the functional importance of these modifications (53). The loss of MeOPN in strain 81-176 increases susceptibility to complement-mediated killing and enhances the invasion of epithelial cells (28, 60), suggesting that MeOPN contributes to CPS-mediated serum resistance and may serve as a key virulence factor in both animal and human hosts (28, 39, 54, 57, 61). These findings support the idea that MeOPN and methylation modifications contribute to immune evasion and host interaction, with phase-variable expression allowing the dynamic adjustment of the cell’s surface architecture. Taken together, our results support the hypothesis that CPS modifications—particularly MeOPN and methylation—function as a modular system.

Specific combinations of ON/OFF states across PVGs fine-tune surface structures to optimize survival under diverse environmental and host-specific pressures. This flexible switching strategy enables *C. jejuni* to navigate immune defenses, antimicrobial stress, and niche-specific colonization.

Previous studies using *C. jejuni* strain NCTC11168 have demonstrated a host-specific enrichment of PVG expression states during mouse colonization, with genes such as *cj1295* and *cj1429* shifting toward the ON phase, and *cj0170* and *cj1306* toward the OFF phase (27, 50, 51, 62). Notably, *CJJ81176_1312* encodes a dimethylglyceric acid modification enzyme homologous to *cj1295*, which has been implicated in flagellar glycosylation and immune evasion (31). Likewise, *CJJ81176_0206* and *CJJ81176_1341* encode homologs of *cj0170* and *cj1306*, respectively, whose OFF-phase enrichment may indicate a selective disadvantage during colonization. These conserved patterns across strains and PVGs reinforce the utility of phase-locked screening platforms like PV-GenShift for dissecting host-specific adaptation mechanisms and identifying functionally relevant PVGs.

PV-GenShift represents a major methodological advancement for investigating PVGs, addressing critical limitations of conventional approaches. Previous studies relied on the use of genetically unstable wild-type populations, typically analyzed as single colonies or pooled isolates, using methods such as Sanger sequencing, fragment length analysis of PCR products, or even multiplex fragment analysis (50, 51, 62, 63). While these approaches have played an important role in characterizing PV, they remain inherently low-throughput and impose substantial logistical burdens, including the need to isolate and analyze large numbers of single colonies to accurately predict PV states while maintaining manageable workloads. Such constraints restrict scalability and introduce variability that complicates causal inference. In contrast, PV-GenShift combines MuGENT-SSR-based genome editing with phase locking, enabling the precise and simultaneous manipulation of multiple loci while eliminating background noise from spontaneous switching.

Rather than focusing solely on detection, this approach actively engineers phase states, stabilizes the genetic context, reduces reliance on labor-intensive workflows, and enhances reproducibility across experimental replicates. By providing a platform readily adaptable to high-throughput functional genomics, PV-GenShift accelerates hypothesis-driven research and opens new avenues for the systematic exploration of PVG function. Despite these advantages, several challenges remain. Editing efficiency did not scale proportionally with the number of MuGENT cycles, and certain loci exhibited persistent OFF-phase bias (Table 2). In addition, this study targeted only 15 PVGs, whereas *Campylobacter* genomes typically harbor a substantially larger repertoire of PVGs (52). Expanding PV-GenShift to accommodate this diversity will require strategies such as iterative multiplex editing, modular donor design, and integration with automated screening pipelines to maintain scalability and phase balance across an extended phasevariome. Addressing these limitations through the optimization of donor DNA design, refinement of transformation protocols, and the development of automated selection strategies will be essential for achieving balanced phasevariomes and fully realizing the scalability of PV-GenShift for comprehensive PVG studies. Ultimately, these advances will position PV-GenShift as a cornerstone technology for dissecting the functional and evolutionary roles of phase variation in bacterial adaptation.

Expanding the platform to include other SSR types (e.g., polyA/T tracts) and non-SSR mechanisms (e.g., promoter inversion or epigenetic phasevarions) will broaden its applicability. The principles of PV-GenShift extend beyond *C. jejuni* to other phase-variable pathogens, including *Neisseria* spp., *Haemophilus influenzae*, and *Helicobacter pylori*. In naturally competent species such as *H. influenzae*, MuGENT-based approaches could be directly implemented (64), while in less transformable bacteria, alternative methods such as CRISPR-based or recombinase-mediated systems may be required. Integrating PV-GenShift with transcriptomic, proteomic, and single-cell analyses will provide deeper insight into PV-regulated networks and phenotypic heterogeneity. Coupling these data with computational modeling and machine learning could enable predictive simulations of phasotype dynamics under varying selective pressures.

In summary, our findings highlight the combinatorial impact of SSR-mediated ON/OFF states on *C. jejuni* adaptation and host interaction. PV-GenShift establishes a generalizable framework for dissecting the roles of PVGs in bacterial pathogenesis. By stabilizing PVG expression and applying defined selective pressures, this platform identifies causative loci underlying immune resistance and colonization, offering a powerful tool for understanding microbial adaptation.

Ultimately, PV-GenShift lays the groundwork for translational applications, including vaccine development, diagnostic marker discovery, and targeted therapeutic design in phase-variable pathogens.

## Materials and Methods

### Bacterial culture conditions

The *C. jejuni* strains used in this study (Table S1) were routinely cultured for 48 hours at 37°C on Brucella broth (BB; Becton Dickinson and Co.) plates containing 1.5% agar (BBA; Kyokuto Pharmaceutical Industrial Co., Ltd.) under microaerophilic conditions using Anaeropack MicroAero gas generator packs (Mitsubishi Gas Chemical Company, Inc.). Motility was assessed by culturing the bacteria on BB plates containing 0.3% agar (BBAM). For liquid culture, fresh single colonies grown on BBA plates were inoculated into 5 mL of Brain Heart Infusion broth (BHB; Becton Dickinson and Co.) in 25 mL test tubes and incubated overnight at 37°C with reciprocal shaking at 160 rpm using a Precyto MG-71M-A Obligatory Anaerobe Culture System (Taitec Corporation). This system efficiently generates a microaerophilic environment by actively aerating gas into each test tube. The gas composition was maintained at 5% O₂, 10% CO₂, and 85% N₂ at a constant flow rate of 10 mL/min. Bacterial stocks were preserved at –80°C in BHB supplemented with 20% glycerol (BHBS). When required, antibiotics were added at the following concentrations: 25 μg/mL chloramphenicol and 50 μg/mL kanamycin.

### DNA manipulation

PCR amplification was carried out using a LifeECO thermal cycler (version 1.04; Bioer Technology Japan Co., Ltd.) with either Quick Taq HS DyeMix DNA polymerase or KOD One PCR Master Mix -Blue (both from Toyobo Co., Ltd.). Sanger sequencing was outsourced to Fasmac Co., Ltd. PCR products were purified using a High Pure PCR Product Purification Kit (Roche Diagnostics K.K.). Custom oligonucleotide primers (Table S2) were obtained from Fasmac Co., Ltd and Hokkaido System Science Co., Ltd. Genomic and plasmid DNA were extracted using a DNeasy Blood and Tissue Kit (Qiagen K.K.) and a High Pure Plasmid Isolation Kit (Roche Diagnostics K.K.), respectively.

### Natural transformation of *C. jejuni* cells using methylated DNA

Natural transformation of *C. jejuni* cells was carried out using methylated donor DNA, following a previously described protocol (53). The procedure consisted of four main steps. First, donor DNA fragments containing specific methylation sites were amplified via PCR (Table S2). Next, the amplified DNA was methylated in *vitro*. The methylated DNA was then introduced into *C. jejuni* cells via natural transformation. Finally, transformants were selected on BBA plates supplemented with the appropriate antibiotics. The specific combinations of template DNA and primer pairs used for PCR amplification are listed in Table S3.

### WGS

WGS was outsourced to Rhelixa, Inc. and performed on an Illumina NovaSeq 6000 platform. Libraries were prepared using a NEBNext Ultra II DNA Library Prep Kit (New England Biolabs Inc.), and sequencing was conducted using a 150 bp paired-end, single-index read format, generating approximately 1 GB of data per sample.

### MuGENT-SSR

Phase-locked *C. jejuni* strains were generated using MuGENT-SSR with methylated donor DNA, as previously described (53). This method utilized both selected and unselected DNA fragments. Selected DNA (e.g., Δ*flaA*::*kan* or Δ*flaA*::*cat*; see Table S2) introduced an antibiotic resistance gene into the *flaA* locus, while unselected DNA introduced locked-ON or locked-OFF mutations into any or all of the 15 PVGs in the *C. jejuni* 81-176 genome (Table S2). Following transformation with the DNA mixture, bacteria were plated on BBA containing the appropriate antibiotic. Forty-eight antibiotic-resistant colonies were cultured overnight in 100 μL of BHB in a 96-well plate and screened for genome editing using MASC PCR. Each 20 μL PCR reaction contained 2 μL of the bacterial culture, 2 μL of a 2.5 μM primer mix (one of six mixes: Mix ON1–3 or Mix OFF1–3; see Table S4), 6 μL of H₂O, and 10 μL of Quick Taq HS DyeMix. The thermal cycling conditions were: 94°C for 5 min and 35 cycles of 94°C for 30 s, 62.6°C for 30 s, and 68°C for 3 min. To eliminate unedited clones, single colonies from positive wells were re-isolated and re-evaluated using MASC PCR.

### Construction of a phase-locked variant library using MuGENT-SSR

A phase-locked variant library was constructed using MuGENT-SSR. The SYC2-0K strain, in which all 15 PVGs were locked in the OFF phase, was cultured in BHB until the A₆₀₀ reached 0.15. One milliliter of this culture was mixed with 0.5 μg of selected DNA (Δ*flaA*::*cat*) and 2 μg of each unselected DNA carrying locked-ON mutations, then incubated at 37°C for 18 h. The resulting chloramphenicol (Cm)-resistant transformants (∼20,000 CFUs) were suspended in 10 mL of BHBS, and genomic DNA was extracted from 1 mL of this suspension.

The remaining suspension was diluted in 5 mL of BHB to an A₆₀₀ of 0.05 and cultured until the A₆₀₀ reached 0.15. A second round of MuGENT-SSR was then performed using selected DNA with a different antibiotic resistance marker (Δ*flaA*::*kan*) and the same unselected DNA. This process was repeated for five cycles, each using a different antibiotic resistance marker. Genomic DNA was extracted from the resulting kanamycin (Km)-resistant transformants and used for subsequent rounds of MuGENT-SSR.

WGS was performed on the extracted DNA, and the number of reads corresponding to locked-ON and locked-OFF mutations in each PVG was quantified using an in-house BLAST+-based tool (65), named “PVfinder_81176.” After trimming raw FASTQ reads with Sickle (66), the reads with 100% identity to either the ON or OFF phase sequences (Table S5) were counted. For each PVG, the ON/OFF ratio was calculated by dividing the number of ON reads by the number of OFF reads. A ratio close to 1 indicates a balanced population (high diversity), while ratios approaching 0 or infinity indicate dominance of OFF or ON variants, respectively. A population with ON/OFF ratios near 1 for all PVGs was selected as a phase-locked library candidate (PLL_81176).

### Restoration of motility in phase-locked variants

Since the *flaA* gene was disrupted by the insertion of an antibiotic resistance cassette during MuGENT-SSR, the resulting phase-locked variants lost motility, a key factor in *C. jejuni* pathogenicity (36, 37, 40, 67). To restore motility, the wild-type *flaA⁺* gene was reintroduced via natural transformation. Approximately 20,000 phase-locked variant colonies from PLL_81176 were suspended in 10 mL of BHB, diluted to an A₆₀₀ of 0.05 in 5 mL of fresh BHB, and cultured until the A₆₀₀ reached 0.15. Transformation was then performed using *flaA⁺* DNA (Table S2).

Briefly, a 50 μL aliquot of the transformation mixture was spotted onto BBAM plates and incubated at 37°C for 24–48 hours. Agar plugs containing motile colonies were then transferred to 5 mL of BHB in a 6-well plate and incubated at 37°C for 24 hours. After centrifugation, the bacterial pellet was resuspended in BHBS to an A₆₀₀ of 0.5. Aliquots (100 μL) of this suspension were stored at –80°C as the final phase-locked variant library, designated PLL_81176M.

### Serum resistance assays

Serum resistance assays were performed based on the method described by Maue et al. (68), with slight modifications. Briefly, 1 mL of an overnight *C. jejuni* culture in 5 mL of BHB was washed twice with 1 mL of minimal essential medium (MEM; Nacalai Tesque, Inc.) and adjusted to an A₆₀₀ of 0.1 in MEM (defined as the initial input bacteria). A 100 μL aliquot of a 1:10,000 dilution of this suspension, containing approximately 5,000 to 10,000 CFUs, was added to each well of a 48-well plate containing 900 μL of prewarmed MEM supplemented with 11.25% human complement serum (HCS; Sigma-Aldrich). The plate was incubated under microaerobic conditions at 37°C for 60 minutes. The percentage of surviving output bacteria after 60 minutes relative to the initial input was determined via serial dilution and plating on mCCDA selective agar (Kanto Chemical Co., Inc.).

### PV_GenShift of human serum-resistant phase variants from PLL_81176M

To select serum-resistant phase variants, the PLL_81176M library was serially passaged in HCS. After each passage, surviving bacteria (1,000–5,000 CFUs) were collected and used as the input for the next round. This process was repeated three times. gDNA was extracted from both the initial input and surviving output bacteria at each passage and subjected to WGS. The ON/OFF ratios of all PVGs were calculated using the PVfinder_81176 tool.

### PV_GenShift of mouse intestinal colonized phase variants from PLL_81176M

Mouse colonization experiments were conducted based on the protocol by Mansfield et al. (57), with slight modifications. C57BL/6 IL-10⁺/⁺ and congenic IL-10⁻/⁻ mice, confirmed to be *Campylobacter*-free, were obtained from The Jackson Laboratory (USA) and maintained under specific pathogen-free conditions at Kyorin University. The PLL_81176M library stored at – 80°C was cultured on tryptic soy agar plates containing 5% sheep blood (Eiken Chemical Co., Ltd.) to yield up to 2,000 colonies per plate. A total of 20,000 colonies were suspended in 5–10 mL of phosphate-buffered saline (PBS) to prepare the input bacterial suspension and extract gDNA. Each mouse (n = 3) was orally inoculated with 0.2 mL of the suspension (3.0 × 10⁸ CFUs) using a sterile 3.5-Fr or 5-Fr feeding tube (Kendall Sovereign; Tyco Healthcare) attached to a 1-mL Luer-Lok syringe (Becton Dickinson). After seven days, the clinical symptoms were assessed, and fecal and cecal samples were collected. Fecal (20–70 mg) and cecal (70–200 mg) contents were suspended in 1 mL of PBS and plated on mCCDA agar to enumerate surviving output colonies. Between 1,000 and 5,000 colonies were pooled in 2 mL of PBS for gDNA extraction. WGS was performed on both input and output samples to determine the ON/OFF ratios of all PVGs. Meanwhile, the wild-type and IL-10-knockout mice were also infected with the parent strain 81-176 (1.0 × 10¹⁰ CFU) as a control.

All experimental protocols were approved by the Biosafety Committee and Animal Ethical Committee at Kyorin University School of Medicine as No.248 (2022/4/19) and No. 257 (2022/8/1), respectively, and conducted in accordance with Guidelines for the Proper Conduct of Animal Experiments (Scientific Council of Japan, 2006/6/1).

### PV_GenShift of chicken intestinal colonized phase variants from PLL_81176M

Chicken colonization experiments were conducted based on the method described by Jones et al. (64), with slight modifications. The PLL_81176M library was cultured on BBA plates to yield up to 2,000 colonies per plate. A total of 20,000 colonies were suspended in 5–10 mL of PBS to prepare the input bacterial suspension and extract the gDNA. Approximately 10⁴ CFUs were orally administered to individually housed, *Campylobacter*-negative Hyline Brown chickens (n = 3) at 27 days of age. Nineteen days post-inoculation, the cecal droppings were collected, suspended in PBS, and plated on BBA supplemented with CCDA selective supplement (Kanto Chemical Co., Inc.) (BBAS). mCCDA was not used for plating bacterial suspensions from chickens, as colonies grown on this medium were occasionally difficult to recover. Between 1,000 and 5,000 colonies of the surviving output bacteria were pooled in 2 mL of PBS for gDNA extraction. WGS was performed on both input and output samples to calculate the ON/OFF ratios of all PVGs.

All animal experiments were conducted in accordance with the animal experiment rules set out in the Animal Welfare Law and Guide for the Care and Use of Laboratory Animals in Obihiro University of Agriculture and Veterinary Medicine (OUAVM). All efforts were made to minimize the suffering of the animals during the experiments. Fertilized eggs were purchased from a commercial hatchery and hatched under the strictly controlled hygienic facility in OUAVM, and the chicks were housed and provided water and food *ad libitum*. The animal experimentation protocol was approved by the President of OUAVM through the oversight of the Institutional Animal Care and Use Committee of the OUAVM (Approval No. 23-180).

## Acknowledgments

This work was supported by the Food Safety Commission of Japan through the Research Program for Risk Assessment Study on Food Safety (Grant Number JPCAFSC20222205), and by the Japan Agency for Medical Research and Development (Grant Numbers JP24fk0108702j0101 and 25fk0108702h0002).

