## Supporting Information for "Genome-scale dissection of phase-variable gene function in *Campylobacter jejuni* using a stabilized phasotype library"

### Figures

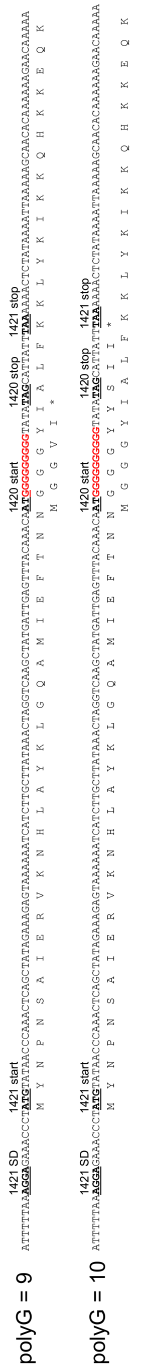

**Fig. S1. Genomic organization and reading frames of *CJJ81176_1421* and *CJJ81176_1420*.** This figure illustrates the relative genomic positions and reading frames of the adjacent genes *CJJ81176_1421* and *CJJ81176_1420* in *Campylobacter jejuni* strain 81-176. The polyG tract (highlighted in red) varies in length (e.g., 9 or 10 G residues), resulting in a frameshift that alters the downstream coding sequence. The start and stop codons for both genes are indicated. Notably, a clear Shine–Dalgarno (SD) sequence is present upstream of *CJJ81176_1421*, but no such sequence is evident for *CJJ81176_1420*. Given the shared reading frame and the absence of an independent SD sequence for *CJJ81176_1420*, this gene is likely a misannotation. These observations suggest that this region represents a single phase-variable gene, *CJJ81176_1421*, whose expression is regulated by changes in the length of the SSR tract.

**Tables**

**Table S1.** Strains and plasmids used in this study.

| Strain or plasmid | Characteristics | Source or reference |
|---|---|---|
| Strain | | |
| 81-176 | *C. jejuni* raw milk origin, serotype R (antigenic factors HS23/HS36) | (1) |
| SYC2-0K | 81-176 Δ*flaA*::*kan* *CJJ81176_0086*$^{OFF}$ *CJJ81176_0206*$^{OFF}$ *CJJ81176_0646*$^{OFF}$ *CJJ81176_0708*$^{OFF}$ *CJJ81176_0758*$^{OFF}$ *CJJ81176_1160*$^{OFF}$ *CJJ81176_1312*$^{OFF}$ *CJJ81176_1325*$^{OFF}$ *CJJ81176_1327*$^{OFF}$ *CJJ81176_1341*$^{OFF}$ *CJJ81176_1419*$^{OFF}$ *CJJ81176_1421*$^{OFF}$ *CJJ81176_1429*$^{OFF}$ *CJJ81176_1432*$^{OFF}$ *CJJ81176_1435*$^{OFF}$ (PT0) | This study |
| SYC2-0 | SYC2-0K *flaA*$^{+}$ | This study |
| SYC2-SV1 | A PLL_81176 variant resistant to human complement serum *CJJ81176_0086*$^{OFF}$ *CJJ81176_0206*$^{ON}$ *CJJ81176_0646*$^{OFF}$ *CJJ81176_0708*$^{OFF}$ *CJJ81176_0758*$^{ON}$ *CJJ81176_1160*$^{OFF}$ *CJJ81176_1312*$^{OFF}$ *CJJ81176_1325*$^{ON}$ *CJJ81176_1327*$^{OFF}$ *CJJ81176_1341*$^{ON}$ *CJJ81176_1419*$^{ON}$ *CJJ81176_1421*$^{OFF}$ *CJJ81176_1429*$^{ON}$ *CJJ81176_1432*$^{OFF}$ *CJJ81176_1435*$^{ON}$ (PT9397) | This study |
| SYC2-SV2C | SYC2-SV1 Δ*flaA*::*cat* *CJJ81176_0086*$^{OFF}$ *CJJ81176_0206*$^{OFF}$ *CJJ81176_0646*$^{OFF}$ *CJJ81176_0708*$^{OFF}$ *CJJ81176_0758*$^{OFF}$ *CJJ81176_1160*$^{OFF}$ *CJJ81176_1312*$^{OFF}$ *CJJ81176_1325*$^{OFF}$ *CJJ81176_1327*$^{OFF}$ *CJJ81176_1341*$^{OFF}$ *CJJ81176_1419*$^{ON}$ *CJJ81176_1421*$^{OFF}$ *CJJ81176_1429*$^{ON}$ *CJJ81176_1432*$^{OFF}$ *CJJ81176_1435*$^{ON}$ (PT21) | This study |
| SYC2-SV2 | SYC2-SV2C *flaA*$^{+}$ | This study |
| SYC2-SV3 | SYC2-SV2C *flaA*$^{+}$ *CJJ81176_0086*$^{OFF}$ *CJJ81176_0206*$^{OFF}$ *CJJ81176_0646*$^{OFF}$ *CJJ81176_0708*$^{OFF}$ *CJJ81176_0758*$^{OFF}$ *CJJ81176_1160*$^{OFF}$ *CJJ81176_1312*$^{OFF}$ *CJJ81176_1325*$^{OFF}$ *CJJ81176_1327*$^{OFF}$ *CJJ81176_1341*$^{OFF}$ *CJJ81176_1419*$^{OFF}$ *CJJ81176_1421*$^{OFF}$ *CJJ81176_1429*$^{ON}$ *CJJ81176_1432*$^{OFF}$ *CJJ81176_1435*$^{ON}$ (PT5) | This study |
| SYC2-SV4 | SYC2-SV2C *flaA*$^{+}$ *CJJ81176_0086*$^{OFF}$ *CJJ81176_0206*$^{OFF}$ *CJJ81176_0646*$^{OFF}$ *CJJ81176_0708*$^{OFF}$ *CJJ81176_0758*$^{OFF}$ *CJJ81176_1160*$^{OFF}$ *CJJ81176_1312*$^{OFF}$ *CJJ81176_1325*$^{OFF}$ *CJJ81176_1327*$^{OFF}$ *CJJ81176_1341*$^{OFF}$ *CJJ81176_1419*$^{ON}$ *CJJ81176_1421*$^{OFF}$ *CJJ81176_1429*$^{OFF}$ *CJJ81176_1432*$^{OFF}$ *CJJ81176_1435*$^{ON}$ (PT17) | This study |
| SYC2-SV5 | SYC2-SV2C *flaA*$^{+}$ *CJJ81176_0086*$^{OFF}$ *CJJ81176_0206*$^{OFF}$ *CJJ81176_0646*$^{OFF}$ *CJJ81176_0708*$^{OFF}$ *CJJ81176_0758*$^{OFF}$ *CJJ81176_1160*$^{OFF}$ *CJJ81176_1312*$^{OFF}$ *CJJ81176_1325*$^{OFF}$ *CJJ81176_1327*$^{OFF}$ *CJJ81176_1341*$^{OFF}$ *CJJ81176_1419*$^{ON}$ *CJJ81176_1421*$^{OFF}$ *CJJ81176_1429*$^{ON}$ *CJJ81176_1432*$^{OFF}$ *CJJ81176_1435*$^{OFF}$ (PT20) | This study |
| SYC2-SV6 | SYC2-SV2C *flaA*$^{+}$ *CJJ81176_0086*$^{OFF}$ *CJJ81176_0206*$^{OFF}$ *CJJ81176_0646*$^{OFF}$ *CJJ81176_0708*$^{OFF}$ *CJJ81176_0758*$^{OFF}$ *CJJ81176_1160*$^{OFF}$ *CJJ81176_1312*$^{OFF}$ *CJJ81176_1325*$^{OFF}$ *CJJ81176_1327*$^{OFF}$ *CJJ81176_1341*$^{OFF}$ *CJJ81176_1419*$^{OFF}$ *CJJ81176_1421*$^{OFF}$ *CJJ81176_1429*$^{OFF}$ *CJJ81176_1432*$^{OFF}$ *CJJ81176_1435*$^{OFF}$ (PT0) | This study |
| SYC2005 | 81-176 Δ*kpsE*::*kan* | This study |
| Plasmid | | |
| pSYC-*cat* | pUCFa *cat* from *C. coli* | (2) |

| | | |
|---|---|---|
| pSYC-*kan* | pUCFa *kan* from *C. coli* | (2) |

**Table S2.** Primers used in this study.

| Primer name | Sequence (5′ to 3′ direction)* |
|---|---|
| 176_1439-f1E | GGGGAATTCTACCTGGTTAACTCCTCGTC |
| 176_1439-kan-r1 | AATGGTTCGCTGGGTTTATCCTTGGTGCTGCAATCAATGT |
| kan-176_1439-f1 | CCTAGATTTAGATGTCTAAAAAGCGTCCAGATATTCCAGAAAGC |
| 176_1439-r1E | GGGGAATTCCTCAGGGTGAAATTCTACCTC |
| 6_0086-f1E | GGGGAATTCGAGTCGTGAAATGGTGATTTTGTAG |
| 6_0086-ON-f1 | GAAGTGCATTTAACTTGGGGCGGAGTAATAGGCTTTAGGG |
| 6_0086-ON-r1 | CCCTAAAGCCTATTACTCCGCCCCAAGTTAAATGCACTTC |
| 6_0086-OFF(-1)-f1 | GAAGTGCATTTAACTTGGGGAGGCTAATAGGCTTTAGGG |
| 6_0086-OFF(-1)-r1 | CCCTAAAGCCTATTAGCCTCCCCAAGTTAAATGCACTTC |
| 6_0086-MASCR1 | GCGGAGAGAGAAAATAAAGCATC |
| 6_0086-MASCmF2 | GAAGTGCATTTAACTTGGGGAGCC |
| 6_0086-MASCmF1 | GAAGTGCATTTAACTTGGGGCGCA |
| 6_0086-r1E | GGGGAATTCGTCTAGCTTTTGTGATCTTCCC |
| 6_0206-f1E | GGGGAATTCGAGCAATGATGCATAAAATGAAGGTG |
| 6_0206-ON-f1 | CTAAGTATTTTAAAAATATAACCGGCGGAGGTATAGAGCCTTATGGC |
| 6_0206-ON-r1 | GCCATAAGGCTCTATACCTCCGCCGGTTATATTTTTAAAATACTTAG |
| 6_0206-OFF(-1)-f1 | CTAAGTATTTTAAAAATATAACCGGAGGCGTATAGAGCCTTATGGC |
| 6_0206-OFF(-1)-r1 | GCCATAAGGCTCTATACGCCTCCGGTTATATTTTTAAAATACTTAG |
| 6_0206-MASCR1 | ACTATAGCCTTTATCGCTCATCATG |
| 6_0206-MASCmF2 | CTAAGTATTTTAAAAATATAACCGGAGCC |
| 6_0206-MASCmF1 | CTAAGTATTTTAAAAATATAACCGGCGCA |
| 6_0206-r1E | GGGGAATTCGAGCAATGATGCATAAAATGAAGGTG |
| 6_0646-f1E | GGGGAATTCGACTCAAAATCTCCTGAAAATTCAGG |
| 6_0646-ON-f1 | CCATTTAAACTAATGAGGGGCGGAGGTATTAGAACGATTTTG |
| 6_0646-ON-r1 | CAAAATCGTTCTAATACCTCCGCCCCTCATTAGTTTAAATGG |
| 6_0646-OFF(-1)-f1 | CCATTTAAACTAATGAGGGGAGGCGTATTAGAACGATTTTG |
| 6_0646-OFF(-1)-r1 | CAAAATCGTTCTAATACGCCTCCCCTCATTAGTTTAAATGG |
| 6_0646-MASCR1 | CATATGTTCCATGATATCTAGTAAAATCG |
| 6_0646-MASCmF2 | CCATTTAAACTAATGAGGGGAGCC |
| 6_0646-MASCmF1 | CCATTTAAACTAATGAGGGGCGCA |
| 6_0646-r1E | GGGGAATTCGACTCAAAATCTCCTGAAAATTCAGG |
| 6_0708-f1E | GGGGAATTCAACCTCATCTTCAACTTCGGC |
| 6_0708-ON-f1 | CTTGCTATAAATTTTAATTTTACCCCACCGCATAAAGATAAATTAG |
| 6_0708-ON-r1 | CTAATTTATCTTTATGCGGTGGGGTAAAATTAAAATTTATAGCAAG |
| 6_0708-OFF(-1)-f1 | CTTGCTATAAATTTTAATTTTACCCCGCCAATAAAGATAAATTAG |
| 6_0708-OFF(-1)-r1 | CTAATTTATCTTTATTGGCGGGGTAAAATTAAAATTTATAGCAAG |

| | |
|---|---|
| 6_0708-MASCR1 | CATCATTTTTGATTCTGTTTCATCTATGG |
| 6_0708-MASCmF2 | CTTGCTATAAATTTTAATTTTACCCCGCGA |
| 6_0708-MASCmF1 | CTTGCTATAAATTTTAATTTTACCCCACGG |
| 6_0708-r1E | GGGGAATTCTGACACAGGTAGTGGTAAGAC |
| 6_0758-f1E | GGGGAATTCGAAAGCATAGCCATAAAATGCG |
| 6_0758-ON-f1 | CGTTTACTGACAGGCGGGGCGGAGATTTAACAATCAAACC |
| 6_0758-ON-r1 | GGTTTGATTGTTAAATCTCCGCCCCGCCTGTCAGTAAACG |
| 6_0758-OFF(-1)-f1 | CGTTTACTGACAGGCGGGGAGGCATTTAACAATCAAACC |
| 6_0758-OFF(-1)-r1 | GGTTTGATTGTTAAATGCCTCCCCGCCTGTCAGTAAACG |
| 6_0758-MASCR1 | GTAATGTTCTTCCACCGTAAATTTCTCC |
| 6_0758-MASCmF2 | CGTTTACTGACAGGCGGGGAGCC |
| 6_0758-MASCmF1 | CGTTTACTGACAGGCGGGGCGCA |
| 6_0758-r1E | GGGGGAATTCGCATCATCTGAAACTTCAAAGC |
| 6_1312-f1E | GGGGAATTCGCATGCGTTATTTTATAGGGG |
| 6_1312-ON-f1 | GAAATTTTAAATAAAACTCTGGGCGGAGGTATACTCAAATTTCACTC |
| 6_1312-ON-r1 | GAGTGAAATTTGAGTATACCTCCGCCCAGAGTTTTATTTAAAATTTC |
| 6_1312-OFF(-1)-f1 | GAAATTTTAAATAAAACTCTGGGAGGCGTATACTCAAATTTCACTC |
| 6_1312-OFF(-1)-r1 | GAGTGAAATTTGAGTATACGCCTCCCAGAGTTTTATTTAAAATTTC |
| 6_1312-MASCR1 | CGTTCCATTCATCAGGGACTATC |
| 6_1312-MASCmF2 | GAAATTTTAAATAAAACTCTGGGAGCC |
| 6_1312-r1E | GGGGAATTCACCTAGGTAGACTATTTTTGC |
| 6_1312-MASCmF1 | GAAATTTTAAATAAAACTCTGGGCGCA |
| 6_1325-f1E | GGGGAATTCGGAGTAAGCATAGCTCATAAG |
| 6_1325-ON-f1 | CTTTAAAATTCAAACTTTAGGCGGAGGGTATCACAAAAAATTGGC |
| 6_1325-ON-r1 | GCCAATTTTTTGTGATACCCTCCGCCTAAAGTTTGAATTTTAAAG |
| 6_1325-OFF(-1)-f1 | CTTTAAAATTCAAACTTTAGGAGGCGGTATCACAAAAAATTGGC |
| 6_1325-OFF(-1)-r1 | GCCAATTTTTTGTGATACCCTCCGCCTAAAGTTTGAATTTTAAAG |
| 6_1325-MASCR1 | CCATACTCTCATTGATATCTGTTC |
| 6_1325-MASCmF2 | CTTTAAAATTCAAACTTTAGGAGCC |
| 6_1325-MASCmF1 | CTTTAAAATTCAAACTTTAGGCGCA |
| 6_1325-r3E | GGGGAATTCAGCTCCTTGGTATGTGGATG |
| 6_1327-f1E | GGGGAATTCGCTTTGCGATTGTCTTTGAAAAC |
| 6_1327-ON-f1 | GCAGATTTACCAAAAATTTATGGCGGAGGGTCTTATGGAGGATAC |
| 6_1327-ON-r1 | GTATCCTCCATAAGACCCTCCGCCATAAATTTTTGGTAAATCTGC |
| 6_1327-OFF(-1)-f1 | GCAGATTTACCAAAAATTTATGGAGGCGGTCTTATGGAGGATAC |
| 6_1327-OFF(-1)-r1 | GTATCCTCCATAAGACCGCCTCCATAAATTTTTGGTAAATCTGC |
| 6_1327-MASCR1 | CTATCAGTAATCCTCATGCCATG |
| 6_1327-MASCmF2 | GCAGATTTACCAAAAATTTATGGAGCC |
| 6_1327-MASCmF1 | GCAGATTTACCAAAAATTTATGGCGCA |
| 6_1327-r2E | GGGGAATTCTGGCTTTGCCAAATGAGGGTT |

| | |
|---|---|
| 6_1341-f1E | GGGGAATTCTGAACTTACCGCTGTTATCGC |
| 6_1341-ON-f1 | GCTCCTTTTGATACTATGGGCGGAGGTATACCAGTTATCATGATAGG |
| 6_1341-ON-r1 | CCTATCATGATAACTGGTATACCTCCGCCCATAGTATCAAAAGGAGC |
| 6_1341-OFF(-1)-f1 | GCTCCTTTTGATACTATGGGAGGCGTATACCAGTTATCATGATAGG |
| 6_1341-OFF(-1)-r1 | CCTATCATGATAACTGGTATACGCCTCCCATAGTATCAAAAGGAGC |
| 6_1341-MASCR1 | CATCAGGTGGAGCTATTTCTTTATC |
| 6_1341-MASCmF2 | GCTCCTTTTGATACTATGGGAGCC |
| 6_1341-MASCmF1 | GCTCCTTTTGATACTATGGGCGCA |
| 6_1341-r3E | GGGGAATTCTTTTAGCACCACGATTGGTTG |
| 81176_1160-f1E | GGGGAATTCTAGGTGCGGTTGCACTTGGTG |
| 6_1160-ON-f1 | GGAAATTATGGATTCATAGGCGGAGGGGATCAACTCTTG |
| 6_1160-ON-r1 | CAAGAGTTGATCCCCTCCGCCTATGAATCCATAATTTCC |
| 81176_1160-OFF(-1)-f1 | GGAAATTATGGATTCATAGGAGGCGGGATCAACTCTTG |
| 81176_1160-OFF(-1)-r1 | CAAGAGTTGATCCCGCCTCCTATGAATCCATAATTTCC |
| 81176_1160-MASCR1 | ACAGGGCGTTTTAGCATGGC |
| 81176_1160-MASCmF1 | AATAGTGGAAATTATGGATTCATAGGAGCC |
| 6_1160-MASCmF1 | AATAGTGGAAATTATGGATTCATAGGCGCA |
| 81176_1160-r1E | GGGGAATTCTTGATGCCTTTGTACATAGGG |
| 81176_1419-f1E | GGGGAATTCATTTGTCCTATATGCTGGGTG |
| 6_1419-ON-f1 | CGTATATTGACAGGCGGAGGGTATTTTACCGCGATTTGG |
| 6_1419-ON-r1 | CCAAATCGCGGTAAAATACCCTCCGCCTGTCAATATACG |
| 81176_1419-OFF(-1)-f1 | CGTATATTGACAGGAGGCGGTATTTTACCGCGATTTGG |
| 81176_1419-OFF(-1)-r1 | CCAAATCGCGGTAAAATACCGCCTCCTGTCAATATACG |
| 81176_1419-MASCR1 | TGCCACTGCTTACACGAGCA |
| 81176_1419-MASCmF1 | CCAGAATTTAATCGTATATTGACAGGAGCC |
| 6_1419-MASCmF1 | CCAGAATTTAATCGTATATTGACAGGCGCA |
| 81176_1419-r1E | GGGGAATTCTGGCTTTATCTCCTAAAACCC |
| 81176_1421-f1E | GGGGAATTCCTTTGCATACATACTCAGATG |
| 6_1421-ON-f1 | GCTATGATTGAGTTTACAAACAATGGCGGAGGGTATATAGC |
| 6_1421-ON-r1 | GCTATATACCCTCCGCCATTGTTTGTAAACTCAATCATAGC |
| 6_1421-OFF(-1)-f1 | GCTATGATTGAGTTTACAAACAATGGAGGCGGTATATAGC |
| 6_1421-OFF(-1)-r1 | GCTATATACCGCCTCCATTGTTTGTAAACTCAATCATAGC |
| 81176_1421-MASCR1 | GCCTTGATTAAAACTTCACCCAGCA |
| 6_1421-MASCmF2 | TGATTGAGTTTACAAACAATGGAGCC |
| 6_1421-MASCmF1 | TGATTGAGTTTACAAACAATGGCGCA |
| 81176_1429-f2E | GGGGAATTCAGATGATGGTGTTGCTACAAG |
| 6_1429-ON-f1 | GTAATGTATAATGGCGGAGGGTATATGAGCAATATTG |
| 6_1429-ON-r1 | CAATATTGCTCATATACCCTCCGCCATTATACATTAC |
| 6-1429-OFF(-1)-f1 | GTAATGTATAATGGAGGCGGTATATGAGCAATATTG |
| 6-1429-OFF(-1)-r1 | CAATATTGCTCATATACCGCCTCCATTATACATTAC |
| 81176_1429-MASCR1 | AACCCCATCTTGCTCTTCAGGA |
| 81176_1429-MASCmF2 | GATGGTGGTTATGTAATGTATAATGGAGCC |
| 6_1429-MASCmF1 | GATGGTGGTTATGTAATGTATAATGGCGCA |

| | |
|---|---|
| 81176_1429-r2E | GGGGAATTCGGAGCTCCAACTAAAGCAGCC |
| 81176_1432-f3E | GGGGAATTCTGGATGGGGTTGAAGTGGTC |
| 6_1432-ON-f1 | GCTAGTAAGAATTGGTATGGCGGAGGGTATATCAAGTTGC |
| 6_1432-ON-r1 | GCAACTTGATATACCCTCCGCCATACCAATTCTTACTAGC |
| 81176_1432-OFF(-1)-f1 | GCTAGTAAGAATTGGTATGGAGGCGGTATATCAAGTTGC |
| 81176_1432-OFF(-1)-r1 | GCAACTTGATATACCGCCTCCATACCAATTCTTACTAGC |
| 81176_1432-MASCR2 | AAACTGTAAACTTAAGTCATGTTGGTGTAC |
| 81176_1432-MASCmF1 | ATTAAAGCTAGTAAGAATTGGTATGGAGCC |
| 6_1432-MASCmF1 | ATTAAAGCTAGTAAGAATTGGTATGGCGCA |
| 81176_1432-r2E | GGGGAATTCCATTACATAACCACCATCTCC |
| 81176_1435-f1E | GGGGAATTCAAATTTGGGCGCTGTTTATGC |
| 6_1435-ON-f1 | GCTATGATTGAGTTTACAAACAATGGCGGAGGGTATATAGC |
| 6_1435-ON-r1 | GCTATATACCCTCCGCCATTGTTTGTAAACTCAATCATAGC |
| 81176_1435-OFF(-1)-f1 | GCTATGATTGAGTTTACAAACAATGGAGGCGGTATATAGC |
| 81176_1435-OFF(-1)-r1 | GCTATATACCGCCTCCATTGTTTGTAAACTCAATCATAGC |
| 81176_1435-MASCR3 | TGAAAAGCTTTCTCTCCTGTTCCATG |
| 81176_1435-MASCmF1 | CTATGATTGAGTTTACAAACAATGGAGCC |
| 6_1435-MASCmF1 | CTATGATTGAGTTTACAAACAATGGCGCA |
| 81176_1435-r1E | GGGGAATTCCTTGCTATGACAACAGCCACG |

*Single lines and double lines indicate edited bases and *Eco*RI sites for methylation, respectively.

**Table S2.** Specific combinations of template DNA and primers used to amplify the donor DNA templates.

| First-step PCR | | Second-step PCR | | Generated PCR product |
|---|---|---|---|---|
| Template | Primers | Template | Primers | |
| gDNA 81-176 | 6_0086-f1E and 6_0086-ON-r1 | First-step PCR products | 6_0086-f1E and 6_0086-r1E | *CJJ81176_0086*$^{ON}$ |
| gDNA 81-176 | 6_0086-ON-f1 and 6_0086-r1E | | | |
| gDNA 81-176 | 6_0086-f1E and 6_0086-OFF(-1)-r1 | First-step PCR products | 6_0086-f1E and 6_0086-r1E | *CJJ81176_0086*$^{OFF}$ |
| gDNA 81-176 | 6_0086-OFF(-1)-f1 and 6_0086-r1E | | | |
| gDNA 81-176 | 6_0206-f1E and 6_0206-ON-r1 | First-step PCR products | 6_0206-f1E and 6_0206-r1E | *CJJ81176_0206*$^{ON}$ |
| gDNA 81-176 | 6_0206-ON-f1 and 6_0206-r1E | | | |
| gDNA 81-176 | 6_0206-f1E and 6_0206-OFF(-1)-r1 | First-step PCR products | 6_0206-f1E and 6_0206-r1E | *CJJ81176_0206*$^{OFF}$ |
| gDNA 81-176 | 6_0206-OFF(-1)-f1 and 6_0206-r1E | | | |
| gDNA 81-176 | 6_0646-f1E and 6_0646-ON-r1 | First-step PCR products | 6_0646-f1E and 6_0646-r1E | *CJJ81176_0646*$^{ON}$ |
| gDNA 81-176 | 6_0646-ON-f1 and 6_0646-r1E | | | |
| gDNA 81-176 | 6_0646-f1E and 6_0646-OFF(-1)-r1 | First-step PCR products | 6_0646-f1E and 6_0646-r1E | *CJJ81176_0646*$^{OFF}$ |
| gDNA 81-176 | 6_0646-OFF(-1)-f1 and 6_0646-r1E | | | |
| gDNA 81-176 | 6_0708-f1E and 6_0708-ON-r1 | First-step PCR products | 6_0708-f1E and 6_0708-r1E | *CJJ81176_0708*$^{ON}$ |
| gDNA 81-176 | 6_0708-f1E and 6_0708-r1E | | | |
| gDNA 81-176 | 6_0708-f1E and 6_0708-OFF(-1)-r1 | First-step PCR products | 6_0708-f1E and 6_0708-r1E | *CJJ81176_0708*$^{OFF}$ |
| gDNA 81-176 | 6_0708-OFF(-1)-r1 | | | |

| | | | | |
|---|---|---|---|---|
| | and 6_0708-r1E | | | |
| gDNA 81-176 | 6_0758-f1E and 6_0758-ON-r1 | First-step PCR products | 6_0758-f1E and 6_0758-r1E | *CJJ81176_0758*$^{ON}$ |
| gDNA 81-176 | 6_0758-ON-f1 and 6_0758-r1E | | | |
| gDNA 81-176 | 6_0758-f1E and 6_0758-OFF(-1)-r1 | First-step PCR products | 6_0758-f1E and 6_0758-r1E | *CJJ81176_0758*$^{OFF}$ |
| gDNA 81-176 | 6_0758-OFF(-1)-f1 and 6_0758-r1E | | | |
| gDNA 81-176 | 81176_1160-f1E and 6_1160-ON-r1 | First-step PCR products | 81176_1160-f1E and 81176_1160-r1E | *CJJ81176_1160*$^{ON}$ |
| gDNA 81-176 | cj1437c -ON-f1 and 6_1160-ON-f1 | | | |
| gDNA 81-176 | 81176_1160-f1E and 81176_1160-OFF(-1)-r1 | First-step PCR products | 81176_1160-f1E and 81176_1160-r1E | *CJJ81176_1160*$^{OFF}$ |
| gDNA 81-176 | 81176_1160-OFF(-1)-f1 and 81176_1160-r1E | | | |
| gDNA 81-176 | 6_1312-f1E and 6_1312-ON-r1 | First-step PCR products | 6_1312-f1E and 6_1312-r1E | *CJJ81176_1312*$^{ON}$ |
| gDNA 81-176 | 6_1312-ON-f1 and 6_1312-r1E | | | |
| gDNA 81-176 | 6_1312-f1E and 6_1312-OFF(-1)-r1 | First-step PCR products | 6_1312-f1E and 6_1312-r1E | *CJJ81176_1312*$^{OFF}$ |
| gDNA 81-176 | 6_1312-OFF(-1)-f1 and 6_1312-r1E | | | |
| gDNA 81-176 | 6_1325-f1E and 6_1325-ON-r1 | First-step PCR products | 6_1325-f1E and 6_1325-r3E | *CJJ81176_1325*$^{ON}$ |
| gDNA 81-176 | 6_1325-ON-f1 and 6_1325-r3E | | | |
| gDNA 81-176 | 6_1325-f1E and 6_1325-OFF(-1)-r1 | First-step PCR products | 6_1325-f1E and | *CJJ81176_1325*$^{OFF}$ |

| | | | | |
|---|---|---|---|---|
| gDNA 81-176 | 6_1325-OFF(-1)-f1 and 6_1325-r3E | | 6_1325-r3E | |
| gDNA 81-176 | 6_1327-f1E and 6_1327-ON-r1 | First-step PCR products | 6_1327-f1E and 6_1327-r2E | *CJJ81176_1327*$^{\mathrm{ON}}$ |
| gDNA 81-176 | 6_1327-ON-f1 and 6_1327-r2E | | | |
| gDNA 81-176 | 6_1327-f1E and 6_1327-OFF(-1)-r1 | First-step PCR products | 6_1327-f1E and 6_1327-r2E | *CJJ81176_1327*$^{\mathrm{OFF}}$ |
| gDNA 81-176 | 6_1327-OFF(-1)-f1 and 6_1327-r2E | | | |
| gDNA 81-176 | 6_1341-f1E and 6_1341-ON-r1 | First-step PCR products | 6_1341-f1E and 6_1341-r3E | *CJJ81176_1341*$^{\mathrm{ON}}$ |
| gDNA 81-176 | 6_1341-ON-f1 6_1341-r3E | | | |
| gDNA 81-176 | 6_1341-f1E and 6_1341-OFF(-1)-r1 | First-step PCR products | 6_1341-f1E and 6_1341-r3E | *CJJ81176_1341*$^{\mathrm{OFF}}$ |
| gDNA 81-176 | 6_1341-OFF(-1)-f1 and 6_1341-r3E | | | |
| gDNA 81-176 | 81176_1419-f1E and 6_1419-ON-r1 | First-step PCR products | 81176_1419-f1E and 81176_1419-r1E | *CJJ81176_1419*$^{\mathrm{ON}}$ |
| gDNA 81-176 | 6_1419-ON-f1 and 81176_1419-r1E | | | |
| gDNA 81-176 | 81176_1419-f1E and 81176_1419-OFF(-1)-r1 | First-step PCR products | 81176_1419-f1E and 81176_1419-r1E | *CJJ81176_1419*$^{\mathrm{OFF}}$ |
| gDNA 81-176 | 81176_1419-OFF(-1)-f1 and 81176_1419-r1E | | | |
| gDNA 81-176 | 81176_1421-f1E and 6_1421-ON-r1 | First-step PCR products | 81176_1421-f1E and 81176_1421-r1E | *CJJ81176_1421*$^{\mathrm{ON}}$ |
| gDNA 81-176 | 6_1421-ON-f1 and | | | |

| | | | | |
|---|---|---|---|---|
| | 81176_1421-r1E | | | |
| gDNA 81-176 | 81176_1421-f1E and 6_1421-OFF(-1)-r1 | First-step PCR products | 81176_1421-f1E and 81176_1421-r1E | *CJJ81176_1421*$^{OFF}$ |
| gDNA 81-176 | 6_1421-OFF(-1)-f1 and 81176_1421-r1E | | | |
| gDNA 81-176 | 81176_1429-f2E and 6_1429-ON-r1 | First-step PCR products | 81176_1429-f2E and 81176_1429-r2E | *CJJ81176_1429*$^{ON}$ |
| gDNA 81-176 | 6_1429-ON-f1 and 81176_1429-r2E | | | |
| gDNA 81-176 | 81176_1429-f2E and 6-1429-OFF(-1)-r1 | First-step PCR products | 81176_1429-f2E and 81176_1429-r2E | *CJJ81176_1429*$^{OFF}$ |
| gDNA 81-176 | 6-1429-OFF(-1)-f1 and 81176_1429-r2E | | | |
| gDNA 81-176 | 81176_1432-f3E and 6_1432-ON-r1 | First-step PCR products | 81176_1432-f3E and 81176_1432-r2E | *CJJ81176_1432*$^{ON}$ |
| gDNA 81-176 | 6_1432-ON-f1 and 81176_1432-r2E | | | |
| gDNA 81-176 | 81176_1432-f3E and 81176_1432-OFF(-1)-r1 | First-step PCR products | 81176_1432-f3E and 81176_1432-r2E | *CJJ81176_1432*$^{OFF}$ |
| gDNA 81-176 | 81176_1432-OFF(-1)-f1 and 81176_1432-r2E | | | |
| gDNA 81-176 | 81176_1435-f1E and 6_1435-ON-r1 | First-step PCR products | 81176_1435-f1E and 81176_1435-r1E | *CJJ81176_1435*$^{ON}$ |
| gDNA 81-176 | 6_1435-ON-f1 and 81176_1435-r1E | | | |
| gDNA 81-176 | 81176_1435-f1E and | | 81176_1435-f1E | *CJJ81176_1435*$^{OFF}$ |

| | | | | |
|---|---|---|---|---|
| | 81176_1435-OFF(-1)-r1 | | and 81176_1435-r1E | |
| gDNA 81-176 | 81176_1435-OFF(-1)-f1 and 81176_1435-r1E | First-step PCR products | | |
| gDNA 81-176 | cjj81176_1339-f1E and flaA81176-cat-r1 | First-step PCR products | cjj81176_1339-f1E and cjj81176_1339-r1E | Δ*flaA*::*cat* |
| pSYC-*cat* | c-cat-f1 and c-cat-r2 | | | |
| gDNA 81-176 | cat-flaA81176-f2 and cjj81176_1339-r1E | | | |
| gDNA 81-176 | cjj81176_1339-f1E and flaA81176-kan-f1 | First-step PCR products | cjj81176_1339-f1E and cjj81176_1339-r1E | Δ*flaA*::*kan* |
| pSYC-*kan* | c-kan-f1 and c-kan-r1 | | | |
| gDNA 81-176 | kan-flaA81176-f1 and cjj81176_1339-r1E | | | |
| gDNA 81-176 | cjj81176_1339-f1E and cjj81176_1339-r1E | | | *flaA*+ |
| gDNA 81-176 | 176_1439-f1E and 176_1439-kan-r1 | First-step PCR products | and | Δ*kpsE*::*kan* |
| pSYC-*kan* | c-kan-f1 and c-kan-r1 | | | |
| gDNA 81-176 | kan-176_1439-f1 and 176_1439-r1E | | | |

**Table S3.** Specific combinations of donor DNA molecules and recipient strains used for natural transformation.

| Donor DNA (PCR fragment) | Recipient strain | Resulting strain |
|---|---|---|
| Δ*flaA*::*cat* or Δ*flaA*::*kan*, *CJJ81176_0086*$^{OFF}$, *CJJ81176_0206*$^{OFF}$, *CJJ81176_0646*$^{OFF}$, *CJJ81176_0708*$^{OFF}$, *CJJ81176_0758*$^{OFF}$, *CJJ81176_1160*$^{OFF}$, *CJJ81176_1312*$^{OFF}$, *CJJ81176_1325*$^{OFF}$, *CJJ81176_1327*$^{OFF}$, *CJJ81176_1341*$^{OFF}$, *CJJ81176_1419*$^{OFF}$, *CJJ81176_1421*$^{OFF}$, *CJJ81176_1429*$^{OFF}$, *CJJ81176_1432*$^{OFF}$, *CJJ81176_1435*$^{OFF}$ | 81-176 | SYC2-0K |
| *flaA*$^{+}$ | SYC2-0K | SYC2-0 |
| Δ*flaA*::*cat*, *CJJ81176_0206*$^{OFF}$, *CJJ81176_1325*$^{OFF}$, *CJJ81176_1341*$^{OFF}$ | SYC2-SV1 | SYC2-SV2C |
| *flaA*$^{+}$ | SYC2-SV2C | SYC2-SV2 |
| *flaA*$^{+}$, *CJJ81176_1419*$^{OFF}$ | SYC2-SV2C | SYC2-SV3 |
| *flaA*$^{+}$, *CJJ81176_1429*$^{OFF}$ | SYC2-SV2C | SYC2-SV4 |
| *flaA*$^{+}$, *CJJ81176_1435*$^{OFF}$ | SYC2-SV2C | SYC2-SV5 |
| *flaA*$^{+}$, *CJJ81176_0206*$^{OFF}$, *CJJ81176_1325*$^{OFF}$, *CJJ81176_1341*$^{OFF}$, *CJJ81176_1419*$^{OFF}$, *CJJ81176_1429*$^{OFF}$, *CJJ81176_1435*$^{OFF}$ | SYC2-SV2C | SYC2-SV6 |
| Δ*kpsE*::*kan* | 81-176 | SYC2005 |

**Table S4.** Primer mixes used for MASC PCR.

| Primer pair | Product size (bp) | ON/OFF Phase detected |
|---|---|---|
| Mix ON1 | | |
| 6_0758-MASCmF1 and 6_0758-MASCR1 | 540 | *CJJ81176_0758*$^{ON}$ |
| 6_0708-MASCmF1 and 6_0708-MASCR1 | 417 | *CJJ81176_0708*$^{ON}$ |
| 6_0206-MASCmF1 and 6_0206-MASCR1 | 300 | *CJJ81176_0206*$^{ON}$ |
| 6_0646-MASCmF1 and 6_0646-MASCR1 | 210 | *CJJ81176_0646*$^{ON}$ |
| 6_0086-MASCmF1 and 6_0086-MASCR1 | 104 | *CJJ81176_0086*$^{ON}$ |
| Mix ON2 | | |
| 6_1327-MASCmF1 and 6_1327-MASCR1 | 506 | *CJJ81176_1327*$^{ON}$ |
| 6_1341-MASCmF1 and 6_1341-MASCR1 | 412 | *CJJ81176_1341*$^{ON}$ |
| 6_1325-MASCmF1 and 6_1325-MASCR1 | 322 | *CJJ81176_1325*$^{ON}$ |
| 6_1312-MASCmF1 and 6_1312-MASCR1 | 100 | *CJJ81176_1312*$^{ON}$ |
| Mix ON3 | | |
| 6_1435-MASCmF1 and 81176_1435-MASCR3 | 654 | *CJJ81176_1435*$^{ON}$ |
| 6_1432-MASCmF1 and 81176_1432-MASCR2 | 512 | *CJJ81176_1432*$^{ON}$ |
| 6_1429-MASCmF1 and 81176_1429-MASCR1 | 406 | *CJJ81176_1429*$^{ON}$ |
| 6_1160-MASCmF1 and 81176_1160-MASCR1 | 332 | *CJJ81176_1160*$^{ON}$ |
| 6_1421-MASCmF1 and 81176_1421-MASCR1 | 221 | *CJJ81176_1421*$^{ON}$ |
| 6_1419-MASCmF1 and 81176_1419-MASCR1 | 145 | *CJJ81176_1419*$^{ON}$ |
| Mix OFF1 | | |
| 6_0758-MASCmF2 and 6_0758-MASCR1 | 540 | *CJJ81176_0758*$^{OFF}$ |
| 6_0708-MASCmF2 and 6_0708-MASCR1 | 417 | *CJJ81176_0708*$^{OFF}$ |
| 6_0206-MASCmF2 and 6_0206-MASCR1 | 300 | *CJJ81176_0206*$^{OFF}$ |
| 6_0646-MASCmF2 and 6_0646-MASCR1 | 210 | *CJJ81176_0646*$^{OFF}$ |
| 6_0086-MASCmF2 and 6_0086-MASCR1 | 104 | *CJJ81176_0086*$^{OFF}$ |
| Mix OFF2 | | |
| 6_1327-MASCmF2 and 6_1327-MASCR1 | 506 | *CJJ81176_1327*$^{OFF}$ |
| 6_1341-MASCmF2 and 6_1341-MASCR1 | 412 | *CJJ81176_1341*$^{OFF}$ |

| | | |
|---|---|---|
| 6_1325-MASCmF2 and 6_1325-MASCR1 | 322 | *CJJ81176_1325*$^{OFF}$ |
| 6_1312-MASCmF2 and 6_1312-MASCR1 | 100 | *CJJ81176_1312*$^{OFF}$ |
| Mix OFF3 | | |
| 81176_1435-MASCmF1 and 81176_1435-MASCR3 | 654 | *CJJ81176_1435*$^{OFF}$ |
| 81176_1432-MASCmF1 and 81176_1432-MASCR2 | 512 | *CJJ81176_1432*$^{OFF}$ |
| 81176_1429-MASCmF2 and 81176_1429-MASCR1 | 406 | *CJJ81176_1429*$^{OFF}$ |
| 81176_1160-MASCmF1 and 81176_1160-MASCR1 | 332 | *CJJ81176_1160*$^{OFF}$ |
| 6_1421-MASCmF2 and 81176_1421-MASCR1 | 221 | *CJJ81176_1421*$^{OFF}$ |
| 81176_1419-MASCmF1 and 81176_1419-MASCR1 | 145 | *CJJ81176_1419*$^{OFF}$ |

**Table S5.** The reference sequences used for phasevariome analysis using PVfinder_81176*.

| PVG | Locked-ON sequence | Locked-OFF sequence |
|---|---|---|
| *CJJ81176_0086* | GAAGTGCATTTAACTTGGGG**C**GG**A**GTAATAGGCTTTAGGG | GAAGTGCATTTAACTTGGGG**A**GG**C**TAATAGGCTTTAGGG |
| *CJJ81176_0206* | CTAAGTATTTTAAAAATATAACCGG**C**GG**A**GGTATAGAGCCTTATGGC | CTAAGTATTTTAAAAATATAACCGG**A**GG**C**GTATAGAGCCTTATGGC |
| *CJJ81176_0646* | CCATTTAAACTAATGAGGGG**C**GG**A**GGTATTAGAACGATTTTG | CCATTTAAACTAATGAGGGG**A**GG**C**GTATTAGAACGATTTTG |
| *CJJ81176_0708* | CTTGCTATAAATTTTAATTTTACCCC**A**CC**G**CATAAAGATAAATTAG | CTTGCTATAAATTTTAATTTTACCCC**G**CC**A**ATAAAGATAAATTAG |
| *CJJ81176_0758* | CGTTTACTGACAGGCGGGG**C**GG**A**GATTTAACAATCAAACC | CGTTTACTGACAGGCGGGG**A**GG**C**ATTTAACAATCAAACC |
| *CJJ81176_1160* | GGAAATTATGGATTCATAGG**C**GG**A**GGGGATCAACTCTTG | GGAAATTATGGATTCATAGG**A**GG**C**GGGATCAACTCTTG |
| *CJJ81176_1312* | GAAATTTTAAATAAAACTCTGGG**C**GG**A**GGTATACTCAAATTTCACTC | GAAATTTTAAATAAAACTCTGGG**A**GG**C**GTATACTCAAATTTCACTC |
| *CJJ81176_1325* | CTTTAAAATTCAAACTTTAGG**C**GG**A**GGGTATCACAAAAAATTGGC | CTTTAAAATTCAAACTTTAGG**A**GG**C**GGTATCACAAAAAATTGGC |
| *CJJ81176_1327* | GCAGATTTACCAAAAATTTATGG**C**GG**A**GGGTCTTATGGAGGATAC | GCAGATTTACCAAAAATTTATGG**A**GG**C**GGTCTTATGGAGGATAC |
| *CJJ81176_1341* | GCTCCTTTTGATACTATGGG**C**GG**A**GGTATACCAGTTATCATGATAGG | GCTCCTTTTGATACTATGGG**A**GG**C**GTATACCAGTTATCATGATAGG |
| *CJJ81176_1419* | CGTATATTGACAGG**C**GG**A**GGGTATTTTACCGCGATTTGG | CGTATATTGACAGG**A**GG**C**GGTATTTTACCGCGATTTGG |
| *CJJ81176_1421* | TTTTTAAAGGAGAAACCCTATGTATAACCCAAACTCAGCTATAGAAAGAGTAAAAAATCATCTTGCTTATAAACTAGGTCAAGCTATGATTGAGTTTACAAACAATGG**C**GG**A**GGGTATATAGC | TTTTTAAAGGAGAAACCCTATGTATAACCCAAACTCAGCTATAGAAAGAGTAAAAAATCATCTTGCTTATAAACTAGGTCAAGCTATGATTGAGTTTACAAACAATGG**A**GG**C**GGTATATAGC |
| *CJJ81176_1429* | GTAATGTATAATGG**C**GG**A**GGGTATATGAGCAATATTG | GTAATGTATAATGG**A**GG**C**GGTATATGAGCAATATTG |
| *CJJ81176_1432* | GCTAGTAAGAATTGGTATGG**C**GG**A**GGGTATATCAAGTTGC | GCTAGTAAGAATTGGTATGG**A**GG**C**GGTATATCAAGTTGC |
| *CJJ81176_1435* | AAAAATAAGGAGAAACCCTATGTATAACCCAAACTCAGCTATAGAAAGAGTAAAAAATCATCTTGCTTATAAACTAGGTCAAGCTATGATTGAGTTTACAAACAATGG**C**GG**A**GGGTATATAGC | AAAAATAAGGAGAAACCCTATGTATAACCCAAACTCAGCTATAGAAAGAGTAAAAAATCATCTTGCTTATAAACTAGGTCAAGCTATGATTGAGTTTACAAACAATGG**A**GG**C**GGTATATAGC |

*Edited bases are underlined.

**Table S6.** Raw data of phasevariomes during the construction of the phase-locked library in *Campylobacter jejuni* strain 81-176.

| PVG | ON/OFF Read Counts of PVGs Across MuGENT-SSR Round | | | | | | | | | | | |
|---|---|---|---|---|---|---|---|---|---|---|---|---|
| | 0 | | 1 | | 2 | | 3 | | 4 | | 5 | |
| | OFF | ON | OFF | ON | OFF | ON | OFF | ON | OFF | ON | OFF | ON |
| *CJJ81176_0086* | 1132 | 0 | 673 | 65 | 680 | 84 | 627 | 106 | 613 | 138 | 839 | 225 |
| *CJJ81176_0206* | 907 | 0 | 519 | 106 | 483 | 144 | 426 | 143 | 437 | 174 | 569 | 249 |
| *CJJ81176_0646* | 619 | 0 | 473 | 39 | 357 | 44 | 338 | 54 | 384 | 50 | 465 | 127 |
| *CJJ81176_0708* | 598 | 0 | 425 | 74 | 339 | 61 | 359 | 91 | 372 | 121 | 454 | 172 |
| *CJJ81176_0758* | 677 | 0 | 523 | 47 | 437 | 65 | 412 | 72 | 371 | 79 | 492 | 157 |
| *CJJ81176_1160* | 811 | 0 | 468 | 112 | 368 | 115 | 277 | 253 | 282 | 286 | 283 | 424 |
| *CJJ81176_1312* | 873 | 0 | 575 | 62 | 514 | 53 | 501 | 62 | 461 | 89 | 513 | 180 |
| *CJJ81176_1325* | 808 | 0 | 505 | 99 | 506 | 104 | 426 | 122 | 380 | 208 | 547 | 225 |
| *CJJ81176_1327* | 802 | 0 | 536 | 74 | 482 | 92 | 344 | 117 | 477 | 193 | 552 | 289 |
| *CJJ81176_1341* | 753 | 0 | 574 | 37 | 504 | 62 | 408 | 61 | 468 | 123 | 601 | 140 |
| *CJJ81176_1419* | 866 | 0 | 538 | 82 | 559 | 79 | 360 | 248 | 351 | 216 | 401 | 435 |
| *CJJ81176_1421* | 189 | 0 | 156 | 8 | 132 | 3 | 107 | 7 | 130 | 20 | 135 | 31 |
| *CJJ81176_1429* | 785 | 0 | 440 | 147 | 368 | 175 | 299 | 220 | 206 | 405 | 442 | 410 |
| *CJJ81176_1432* | 883 | 0 | 499 | 136 | 384 | 122 | 387 | 197 | 432 | 229 | 628 | 192 |
| *CJJ81176_1435* | 147 | 0 | 110 | 9 | 82 | 8 | 54 | 30 | 78 | 28 | 79 | 72 |

**Table S7.** Raw data of phasevariomes of human serum-resistant variants from PLL_81176.

| PVG | ON/OFF Read Counts of PVGs | | | | | | | |
|---|---|---|---|---|---|---|---|---|
| | Input | | Output | | | | | |
| | | | 1st | | 2nd | | 3rd | |
| | OFF | ON | OFF | ON | OFF | ON | OFF | ON |
| *CJJ81176_0086* | 840 | 213 | 771 | 129 | 793 | 98 | 814 | 81 |
| *CJJ81176_0206* | 564 | 182 | 515 | 166 | 515 | 180 | 480 | 201 |
| *CJJ81176_0646* | 402 | 123 | 341 | 133 | 440 | 64 | 388 | 35 |
| *CJJ81176_0708* | 383 | 111 | 295 | 111 | 466 | 54 | 453 | 27 |
| *CJJ81176_0758* | 409 | 121 | 236 | 339 | 74 | 483 | 60 | 479 |
| *CJJ81176_1160* | 588 | 33 | 493 | 85 | 592 | 69 | 557 | 33 |
| *CJJ81176_1312* | 569 | 129 | 564 | 122 | 608 | 60 | 642 | 60 |
| *CJJ81176_1325* | 30 | 724 | 70 | 593 | 42 | 663 | 30 | 709 |
| *CJJ81176_1327* | 615 | 116 | 473 | 181 | 685 | 59 | 611 | 50 |
| *CJJ81176_1341* | 27 | 733 | 81 | 625 | 54 | 591 | 27 | 611 |
| *CJJ81176_1419* | 820 | 49 | 139 | 621 | 29 | 714 | 26 | 729 |
| *CJJ81176_1421* | 106 | 8 | 24 | 77 | 24 | 104 | 33 | 90 |
| *CJJ81176_1429* | 78 | 547 | 401 | 184 | 525 | 176 | 464 | 189 |
| *CJJ81176_1432* | 493 | 179 | 335 | 274 | 299 | 429 | 301 | 447 |
| *CJJ81176_1435* | 104 | 13 | 36 | 69 | 9 | 95 | 2 | 113 |

**Table S8.** Raw data of phasevariomes of PLL_81176-derived colonizing variants in the cecum and feces of IL-10-knockout mice.

| PVG | ON/OFF Read Counts of PVGs | | | | | | | | | | | | | |
|---|---|---|---|---|---|---|---|---|---|---|---|---|---|---|
| | | | Output | | | | | | | | | | | |
| | | | m#1 | | | | m#2 | | | | m#3 | | | |
| | Input | | Cecum | | Feces | | Cecum | | Feces | | Cecum | | Feces | |
| | OFF | ON | OFF | ON | OFF | ON | OFF | ON | OFF | ON | OFF | ON | OFF | ON |
| *CJJ81176_0086* | 635 | 185 | 1438 | 0 | 860 | 0 | 1519 | 0 | 1430 | 1 | 722 | 0 | 712 | 0 |
| *CJJ81176_0206* | 414 | 192 | 1102 | 13 | 669 | 6 | 1259 | 9 | 1136 | 10 | 523 | 46 | 468 | 21 |
| *CJJ81176_0646* | 334 | 177 | 717 | 3 | 506 | 6 | 825 | 4 | 887 | 13 | 369 | 4 | 398 | 5 |
| *CJJ81176_0708* | 278 | 198 | 793 | 4 | 497 | 1 | 917 | 0 | 838 | 2 | 334 | 20 | 403 | 11 |
| *CJJ81176_0758* | 403 | 172 | 882 | 0 | 586 | 0 | 1043 | 0 | 919 | 0 | 424 | 0 | 415 | 2 |
| *CJJ81176_1160* | 411 | 173 | 932 | 1 | 565 | 4 | 1054 | 5 | 1032 | 9 | 492 | 4 | 493 | 12 |
| *CJJ81176_1312* | 465 | 207 | 4 | 1074 | 1 | 690 | 2 | 1240 | 1 | 1097 | 38 | 479 | 12 | 510 |
| *CJJ81176_1325* | 357 | 245 | 1083 | 0 | 633 | 0 | 1172 | 0 | 1194 | 0 | 510 | 0 | 541 | 0 |
| *CJJ81176_1327* | 390 | 261 | 1076 | 3 | 649 | 2 | 1108 | 4 | 1111 | 9 | 471 | 7 | 541 | 6 |
| *CJJ81176_1341* | 473 | 166 | 967 | 3 | 562 | 0 | 1165 | 11 | 1105 | 2 | 442 | 1 | 537 | 6 |
| *CJJ81176_1419* | 389 | 300 | 4 | 1287 | 4 | 780 | 2 | 1279 | 1 | 1166 | 42 | 554 | 13 | 568 |
| *CJJ81176_1421* | 80 | 39 | 3 | 203 | 1 | 116 | 2 | 208 | 1 | 224 | 4 | 87 | 2 | 103 |
| *CJJ81176_1429* | 191 | 431 | 5 | 1027 | 2 | 609 | 6 | 1134 | 4 | 1135 | 4 | 520 | 10 | 557 |
| *CJJ81176_1432* | 340 | 319 | 1048 | 0 | 620 | 0 | 1150 | 0 | 1128 | 1 | 481 | 0 | 565 | 4 |
| *CJJ81176_1435* | 107 | 20 | 193 | 0 | 97 | 0 | 219 | 0 | 201 | 4 | 95 | 0 | 93 | 1 |

**Table S9.** Raw data of phasevariome of PLL_81176-derived variants recovered from chicken cecal droppings.

| PVG | ON/OFF Read Counts of PVGs | | | | | | | |
|---|---|---|---|---|---|---|---|---|
| | input | | Output | | | | | |
| | | | c#1 | | c#2 | | c#3 | |
| | OFF | ON | OFF | ON | OFF | ON | OFF | ON |
| *CJJ81176_0086* | 784 | 247 | 466 | 226 | 6 | 1336 | 791 | 0 |
| *CJJ81176_0206* | 535 | 260 | 593 | 0 | 8 | 1127 | 690 | 0 |
| *CJJ81176_0646* | 455 | 259 | 144 | 280 | 1 | 695 | 432 | 0 |
| *CJJ81176_0708* | 411 | 285 | 27 | 369 | 4 | 757 | 495 | 0 |
| *CJJ81176_0758* | 439 | 237 | 482 | 1 | 872 | 0 | 0 | 543 |
| *CJJ81176_1160* | 598 | 221 | 371 | 150 | 5 | 950 | 627 | 0 |
| *CJJ81176_1312* | 558 | 284 | 9 | 630 | 1012 | 5 | 674 | 0 |
| *CJJ81176_1325* | 535 | 279 | 189 | 332 | 0 | 931 | 0 | 592 |
| *CJJ81176_1327* | 542 | 376 | 372 | 151 | 934 | 7 | 0 | 654 |
| *CJJ81176_1341* | 608 | 203 | 582 | 0 | 913 | 0 | 589 | 0 |
| *CJJ81176_1419* | 508 | 287 | 394 | 200 | 3 | 1118 | 0 | 712 |
| *CJJ81176_1421* | 137 | 28 | 23 | 1 | 0 | 193 | 0 | 107 |
| *CJJ81176_1429* | 258 | 563 | 0 | 519 | 0 | 942 | 0 | 611 |
| *CJJ81176_1432* | 446 | 411 | 175 | 331 | 0 | 1021 | 669 | 0 |
| *CJJ81176_1435* | 116 | 62 | 0 | 21 | 1 | 151 | 104 | 0 |
